# Complete phylogeny of *Micrathena* spiders suggests multiple dispersal events among Neotropical rainforests, islands, and landmasses, and indicates Andean orogeny promotes speciation

**DOI:** 10.1101/2023.11.01.565210

**Authors:** Ivan L. F. Magalhaes, Pedro H. Martins, Bárbara T. Faleiro, Teofânia H. D. A. Vidigal, Fabrício R. Santos, Leonardo S. Carvalho, Adalberto J. Santos

## Abstract

The Neotropical region is the most diverse on the planet, largely due to its mosaic of tropical rainforests. Multiple tectonic and climatic processes have been hypothesized to play a role in generating this diversity; these include Andean orogeny, closure of the Panama Isthmus, the hypothesized GAARlandia land bridge, and putative historical connections among currently isolated forests. *Micrathena* spiny spiders (Araneidae) include ∼120 species distributed mostly in forests from Mexico to Argentina, including the Antilles. Here, we use it as a model to study the biogeographic history of Neotropical rainforests by estimating a complete, dated phylogeny using morphological data for all species and molecular data for a subsample of 79 species. This resulted in a phylogeny that is mostly robust and supports most of the previously recognized species groups, although with uncertainty in the phylogenetic position of some species, especially those lacking sequences. The genus began diversifying about 25 million years ago. We use an event-based approach and biogeographic stochastic mapping to estimate ancestral distributions and the timing and direction of dispersal events, and to identify areas where diversity was generated, while accounting for phylogenetic uncertainty. Andean cloud forests generated the majority of species through in-situ speciation, but the Amazon was the major source of species for adjacent areas through dispersal; on the other extreme, the Dry Diagonal received species from other areas but generated very little diversity. Species exchange between Central and South America was intense, with ∼23 dispersal events beginning at least 20 million years ago, indicating that *Micrathena* dispersed between these continents before closure of the Isthmus of Panama. We also inferred ∼4 over-water dispersal events from Central and North America to the Antilles, which happened in the last 20 million years, and thus much after the proposed age of the GAARlandia land bridge. We identified an important species exchange route between the Amazon and the Atlantic forests. Sampling all species of the genus was fundamental to some of the conclusions above, especially in identifying the Andes as the area that generated the majority of species. This study highlights the importance of a solid and complete taxonomic sampling in biogeographic studies.

## Introduction

The Neotropical realm extends from Argentina to Mexico and is the most species-rich region on Earth (Antonelli et al. 2018a). This realm encompasses diverse biomes, ranging from deserts to temperate rainforests. Of special significance are the tropical forests: for instance, the Amazon (Fig. 1, M) is among the most species-rich tropical rainforests in the world (Kier et al. 2005). There is fossil evidence for continued existence of Neotropical rainforests for at least ∼58 million years, with plant species diversity within the same order of magnitude as the extant flora (Wing et al. 2009). Understanding how these ancient and diverse biomes assembled is a major goal of biogeography. Recent syntheses have summarized biogeographic patterns across different taxa by relying on occurrence data and phylogenies (e.g., Antonelli et al. 2015, 2018b, Matos-Maraví et al. 2021). This approach allows identifying areas that generate diversity (through speciation) and the main sources or sinks of species (through dispersal). Some of these works identified the Amazon as a major cradle of diversity that sources species to the surrounding regions (Antonelli et al. 2018b).

**Figure 1.**
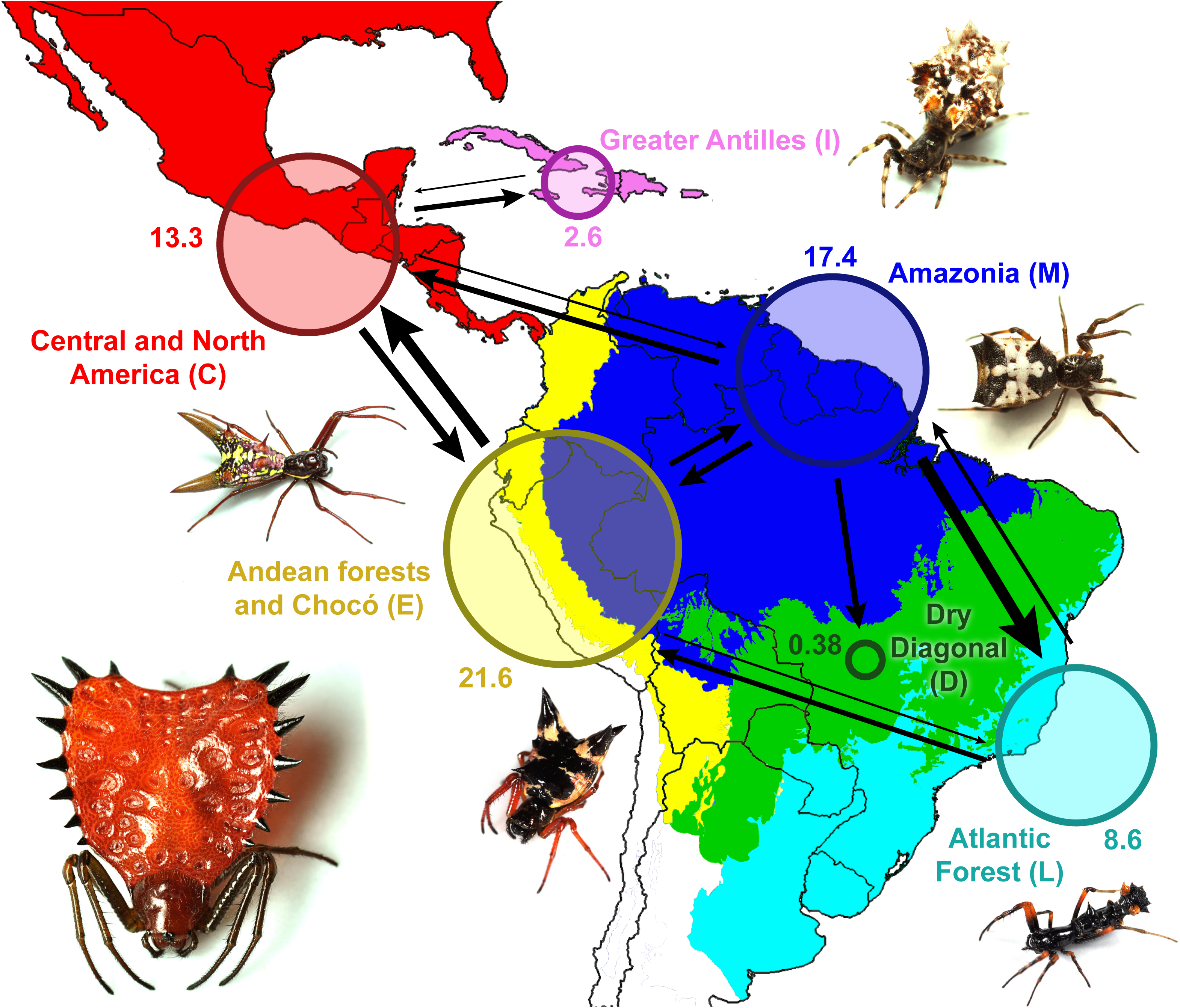
Main centers of origin and dispersal routes of *Micrathena* in the Neotropics. Estimates are based on 10 biogeographic stochastic maps for each of 100 trees from the posterior distribution of the dated analysis. The circles are proportional to the average estimated number of in-situ speciation events in each area. Arrows indicate the direction of dispersal events, with their width proportional to the number of events. Rare dispersal events (mean estimated number of events lower than standard deviation) are not depicted.

The heterogeneity of the Neotropical region provides ample opportunity for testing different biogeographic hypotheses (Antonelli et al. 2018a, see their fig. 1). One of such hypotheses postulates recurring connections among ecologically similar regions that are currently geographically separated. For instance, the Atlantic rainforest, extending from the northeastern coast of Brazil to northeastern Argentina (Fig. 1, L), is one of the world’s biodiversity hotspots (Myers et al. 2000). It is currently isolated from other humid forests (such as the Amazon and Andean cloud forests) by the Dry Diagonal, a corridor of savannahs and dry forests (Fig. 1, D). However, these humid forests have been connected multiple times due to past climatic oscillations. There is evidence of repeated biotic interchange between the Atlantic rainforest and Andean cloud forests, probably starting in the Miocene but extending into the Quaternary (e.g., Trujillo-Arias et al. 2017, 2020). Connections between the Atlantic Forest and the Amazon are better documented, and a recent revision (Ledo & Colli 2017) indicates two main patterns: older Miocene connections between the Amazon and the southern portion of the Atlantic Forest (e.g., Batalha-Filho et al. 2013, Peres et al. 2017, De Sá et al. 2019, Matos-Maraví et al. 2021) and more recent Quaternary connections between the Amazon and the northern portion of the Atlantic Forest (e.g., Prates et al. 2016, Peres et al. 2017, Nascimento et al. 2021, Ledru & Araújo 2023). Humid forests exchanged species not only among themselves, but also with other biomes; for instance, there is evidence for frequent transitions from evergreen rainforests to seasonally dry biomes, with the reverse pattern being less common (Antonelli et al. 2018b, Ziska et al. 2020).

The landscape in the Neotropical region was affected not only by climatic oscillations, but also by tectonic activity. South America has the longest mountain range in the world, the Andes, which extend along the entire western end of the continent (Graham 2009). Andean uplift is the result of the subduction of the Nazca plate beneath the South American plate and began around 40 million years ago (Mya), with some portions of the mountain range attaining half their current height around ∼15–9 Mya (Hartley 2003, Graham 2009). The consequences of Andean uplift for the South American landscape were dramatic, including a change in the course of the proto-Amazon River. The creation of novel habitats promoted the appearance of montane taxa (Hoorn et al. 2010). The complex Andean landscape limits the contact between populations, leading to speciation (e.g., Salgado-Roa et al. 2021, 2022); as a result, topographic barriers often delimit areas of endemism, especially in the tropical portion (Hazzi et al. 2018). There is evidence that Andean uplift promoted diversification in some groups (e.g., Ceccarelli et al. 2016 and references therein), while others display similar diversification rates in montane and lowland clades (e.g., Buianain et al. 2022). In some groups, mountainous regions are associated with higher species richness, but speciation dynamics is similar in mountains and lowlands, and thus the reason for the increased diversity in mountains remains unclear (Vargas et al. 2020).

The biota of South America has evolved in relative isolation for a long time after Gondwana break-up. This isolation has been broken in the Cenozoic by the formation of the Isthmus of Panama, which connects South and Central America (O’Dea et al. 2016), and possibly by hypothetical land bridges uniting South America and the Greater Antilles (GAARlandia; Iturralde-Viñent & MacPhee 1999). The timing of closure of the Isthmus of Panama has been debated. Early studies of marine foraminifera placed this event at ∼3.5 Mya (Saito 1976). This date was generally accepted until new geological (Montes et al. 2012, 2015) and biological (Bacon et al. 2015) data suggested the isthmus may have formed much earlier, in the late Miocene (15–7 Mya). In the most comprehensive revision on the subject, O’Dea et al. (2016) concluded that the existing evidence unambiguously support full closure of the isthmus around 3 Mya. This allowed increased exchange of land organisms between South and Central America (e.g., Matos-Maraví et al. 2021). Nevertheless, fossils and phylogenetic relationships of many groups are consistent with over-water dispersal between these continents preceding the final closure of the isthmus (e.g., Cody et al. 2010, Bloch et al. 2016, Crews & Esposito 2020, Ramos et al. 2022). A similar situation is observed regarding faunal exchanges between continental America and the Antilles. Iturralde-Viñent & MacPhee (1999) put forward the hypothesis of GAARlandia, which postulates that the Greater Antilles were briefly connected to South America through the Aves Ridge between 33–32 Mya. This hypothetical land bridge would have allowed biotic exchange of terrestrial organisms between these areas. The evidence supporting GAARlandia was recently reviewed by Ali & Hedges (2021): they found no geological evidence of emergence of the Aves Ridge during the proposed time of existence of GAARlandia, and no coincidence in the timing of arrival of ∼35 vertebrate groups to the islands. While some groups display phylogenies consistent with GAARlandia (e.g. Schools et al. 2022), others are more consistent with over-water dispersal from the continent to the islands (e.g. Shapiro et al. 2022). Thus, it seems that faunal exchanges between these landmasses (South and Central America, and continental America and the Antilles) has frequently happened in an idiosyncratic manner depending on the dispersal capabilities of each group.

Early comparative works in the Neotropical region already found idiosyncratic biogeographic patterns for different taxa and stated that “given the general scarcity of phylogenies for Neotropical lineages, at this time we should focus on recording specific biogeographical patterns rather than seeking to identify processes of diversification of the whole Neotropical fauna” (Costa 2003). Twenty years later, comprehensive and well-resolved phylogenies are still lacking for many Neotropical groups, especially invertebrates, which precludes complete biogeographic syntheses (but see Crews & Esposito 2020). Most of the studies cited in the paragraphs above rely only in data from vertebrates (e.g., Ledo & Colli 2017, Ali & Hedges 2021) or vertebrates and plants (e.g., Myers et al. 2000, Antonelli et al. 2018b). In the few studies that include invertebrates, they represent only a small fraction of the sampled taxa (e.g., Bacon et al. 2015, Cody et al. 2010), and these acknowledge that their efforts could be improved by the addition of more invertebrates. This is unfortunate, since invertebrates are the major component of animal diversity (Wheeler 1990) and provide important biogeographical insights at all geographic scales (e.g., Peres et al. 2017, Crews & Esposito 2020, Matos-Maraví et al. 2021, Beza-Beza et al. 2021, Ramos et al. 2022). In this context, the need for species-level phylogenies of invertebrates is particularly pressing.

Among invertebrates in Neotropical forests, the spiny spiders in the genus *Micrathena* Sundevall, 1833 (Araneidae) are certainly among the most charismatic. The females of many species are showy and colorful and may bear long abdominal spines (Fig. 2). The genus ranges from Argentina to Canada, including the Antilles, and is comprised by 117 described species, most of which build orb-webs in the understory of tropical forests (WSC 2023). Since the seminal taxonomic revision of the genus by Levi (1985), only three valid species have been described (see Magalhaes & Santos, 2011; Magalhaes et al. 2017), demonstrating that its diversity is well-known. Due to its high diversity and ubiquity in the Neotropical region, *Micrathena* is an excellent model for testing biogeographic hypotheses at multiple scales. However, a comprehensive phylogeny of the genus is lacking; previous efforts are based only on morphological data of selected representatives (Magalhaes & Santos 2012, Magalhaes et al. 2017) or focused on the phylogeography of Caribbean species (McHugh et al. 2014, Shapiro et al. 2022). The latter have suggested that *Micrathena* reached the Antilles via over-water dispersal multiple times, hinting that the genus has decent dispersal capabilities.

**Figure 2.**
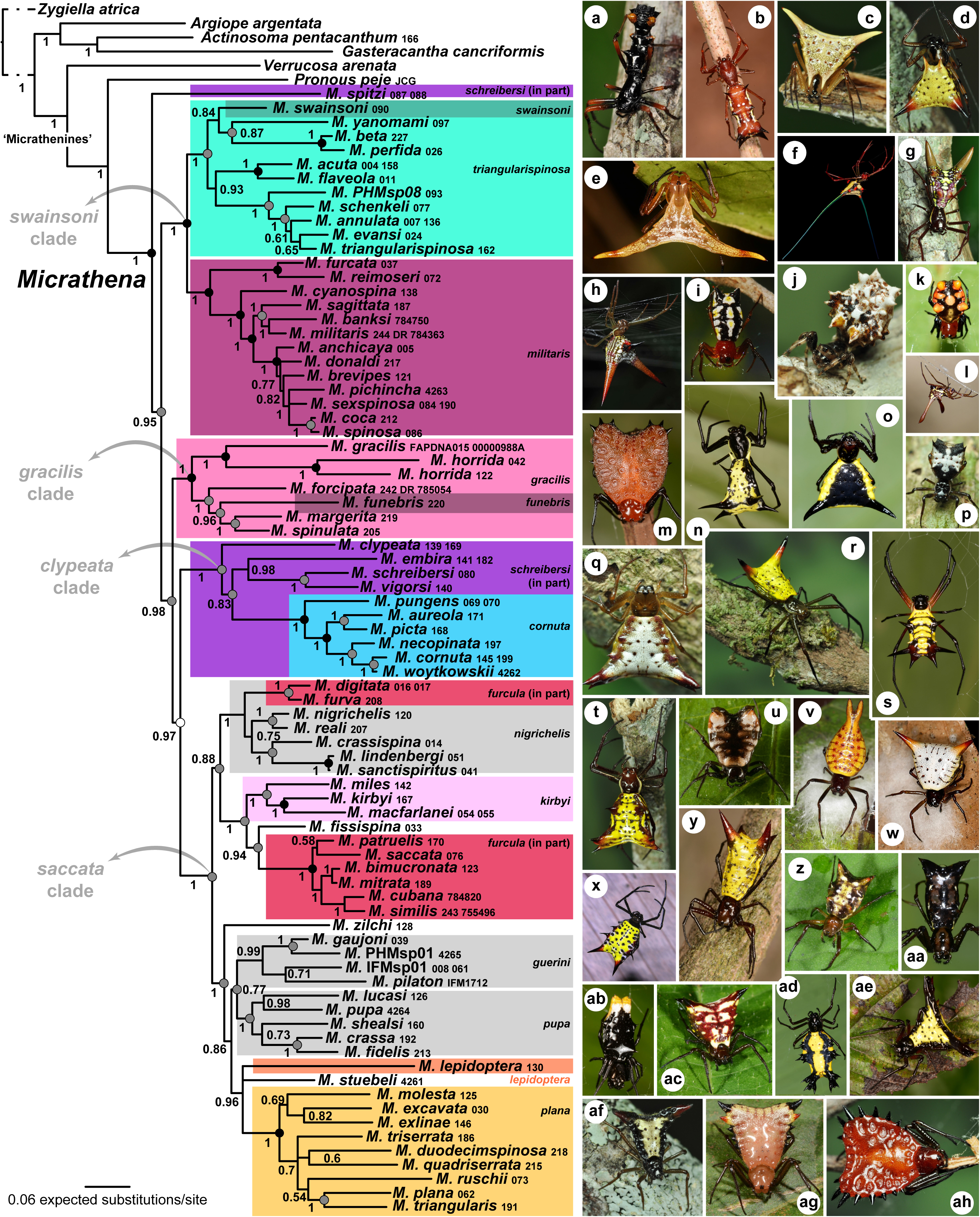
Left panel, phylogenetic relationships among *Micrathena* estimated using Bayesian inference based on data from morphology and five concatenated molecular markers (COI, 16S, H3, 28S, and ITS). Only species for which we sequenced at least one molecular marker are included. Background colors (except gray ones) indicate previously recognized species groups, several of which are not monophyletic; species in the white or gray backgrounds include those previously placed in the *guerini* species group, or *incertae sedis* species. Numbers below branches are posterior probabilities. Nodes with black dots have also been recovered in analyses of datasets containing morphology or sequences only; nodes with grey dots have only been recovered in datasets containing sequences but are contradicted by the morphological dataset; the white dot indicates a node recovered only when morphological data is included. Right panel, photos of live specimens (credits: AA = Arthur Anker, ACA = Alfredo Colón Archilla, ACS = Alexis Callejas Segura, CF = César Favacho, GK = Gernot Kunz, PHM = Pedro H. Martins, RA = Ricardo Arredondo). **a** *Micrathena spitzi* (PHM), **b** *M. swainsoni* (AA & PHM), **c** *M. perfida* (PHM), **d** *M. annulata* (PHM), **e** *M. reimoseri* (AA), **f** *M. cyanospina* (CF), **g** *M. spinosa* (PHM), **h** *M. militaris* (ACA), **i** *M. funebris* (AA), **j** *M. horrida* (PHM), **k** *M. glyptogonoides* (RA), **l** *M. forcipata* (ACS), **m** *M. clypeata* (AA & PHM), **n** *M. vigorsi* (AA & PHM), **o** *M. pungens* (AA & PHM), **p** *M. aureola* (PHM), **q** *M. digitata* (PHM), **r** *M. crassispina* (PHM), **s** *M. kirbyi* (AA & PHM), **t** *M. fissispina* (PHM), **u** *M. patruelis* (PHM), **v** *M. furcula* (GK), **w** *M. bimucronata* (GK), **x** *M. zilchi* (AA), **y** *M. guerini* (AA), **z** *M. fidelis* (AA & PHM), **aa** *M. shealsi* (AA & PHM), **ab** *M. pupa* (AA & PHM), **ac** *M. lucasi* (AA & PHM), **ad** *M. lepidoptera* (AA & PHM), **ae** *M. stuebeli* (AA & PHM), **af** *M. plana* (PHM), **ag** *M. excavata* (AA & PHM), **ah** *M. ruschii* (PHM).

Biogeographic synthesis often requires inferring in-situ speciation and dispersal among areas. Such inferences are usually drawn from estimates of ancestral ranges and biogeographic stochastic mapping (e.g., Xing & Ree 2017, Matos-Maraví et al. 2021, Antonelli et al. 2018b, Magalhaes et al. 2021). However, these methods are very sensitive to the taxon sampling: unsampled lineages are treated as “extinct”, and the existing models consistently underestimate extinction (Ree & Smith 2008), leading to incorrect estimates of ancestral ranges. Thus, complete phylogenies sampling all species within a group are preferrable when taking this approach. However, in many taxa it is difficult to obtain fresh material for sequencing from all species. Some have solved this problem by placing non-sequenced species in the tree by using topological constraints based on morphology (e.g., Landis et al. 2021). We argue that a better alternative would be a total-evidence analysis, where the phylogenetic placement of non-sequenced specimens is estimated, instead of constrained based on prior beliefs. *Micrathena* represents an excellent model to take this approach, since a morphological matrix is already available (Magalhaes et al. 2017) and can be easily expanded to accommodate all species and combined with newly generated sequence data.

The main aims of this paper are: (1) generate a complete, dated, total-evidence phylogenetic hypothesis for *Micrathena* by integrating morphology and molecular data, and use this new systematic framework to test several hypothesis regarding the biogeography of the genus in the Neotropical region; (2) understand the diversification of the genus in a geographic context by identifying which areas are the source of new species through speciation, and the main dispersal routes; (3) estimate the timing and direction of dispersal among Neotropical rainforests that are currently isolated; (4) test whether the dispersal of these spiders to the Greater Antilles is consistent with the GAARlandia land bridge hypothesis; (5) test whether dispersal rates of these spiders between South and Central America increased after the closure of the Panama Isthmus and (6) understand what biogeographic insights can be gained by using a complete phylogeny of the genus, as opposed to a phylogeny including only those species for which fresh material is available for sequencing.

## Material and methods

### Taxon sampling, morphological matrix, and voucher specimens

We aimed at collecting morphological data for all 117 described species of *Micrathena*, as well as three undescribed species examined by us (*Micrathena* IFMsp01 and *Micrathena* PHMsp01 from Ecuador; and *Micrathena* PHMsp08 from Brazil). We could not distinguish specimens of *M. annulata* Reimoser and *M. jundiai* Levi based on the morphological diagnosis of Levi (1985), as the two species intergrade; we suspect they may be synonyms, and all specimens are here treated as *M. annulata*. Thus, our morphological dataset includes 119 *Micrathena* species. Three species formally known only by females (*M. glyptogonoides* Levi, *M. gurupi* Levi and *M. hamifera* Simon) were scored for both sexes based on males found in collections alongside the respective females.

We augmented the morphological matrix of Magalhaes et al. (2017), which included 53 *Micrathena* species, to include the 119 species sampled by us, plus 13 outgroups belonging to other araneid genera. We examined 169 new specimens for morphological scorings. We could not examine specimens of 26 *Micrathena* species directly, and thus their morphological scorings were based on the illustrations by Levi (1985). We revised and re-worked previous scorings and characters as needed and added 18 new characters (resulting in 192 characters in total). No characters were considered ordered.

We obtained sequence data for 188 newly sequenced specimens (11 belonging to outgroups, 177 to *Micrathena*), nearly half of which collected by us in field expeditions. We complemented these with data mined from GenBank and sequences kindly provided by F.M. Labarque and J. Cabra-García (9 outgroups, 45 *Micrathena*). This resulted in a dataset of 222 specimens of 79 species of *Micrathena* and 20 outgroups, sequenced for at least one molecular marker each. We were unable to gather fresh material for 40 species of *Micrathena*, and thus these are represented only in the morphological partition. Outgroups contain a variety of araneid genera, including two Micratheninae found to be successive sister groups to *Micrathena* (*Pronous* and *Verrucosa*; see Cabra-García & Hormiga 2020). Trees were rooted on *Zygiella*.

Voucher specimens used for morphology and sequencing are deposited at these collections: AMNH—American Museum of Natural History (USA); CAS—California Academy of Sciences (USA); CHNUFPI—Coleção de História Natural da Universidade Federal do Piauí (Brazil); CNAN-Ar—Colección Nacional de Arácnidos, Universidad Nacional Autónoma de México (Mexico); IBSP—Instituto Butantan (Brazil); MACN-Ar— Museo Argentino de Ciencias Naturales ‘Bernardino Rivadavia’ (Argentina); MCZ— Museum of Comparative Zoology, Harvard University (USA); MPEG—Museu Paraense Emílio Goeldi (Brazil); MUSM—Museo de Historia Natural de la Universidad Nacional Mayor de San Marcos (Peru); MZSP—Museu de Zoologia da Universidade de São Paulo (Brazil); UFMG—Centro de Coleções Taxonômicas da Universidade Federal de Minas Gerais (Brazil).

### DNA sequencing

We extracted genomic DNA of specimens preserved at-20 °C in 96% ethanol, or specimens in 80% ethanol stored at room temperature for no more than 8 years at the time of extraction. We extracted genomic DNA of 1–4 legs per individual using the Promega Wizard^®^ Genomic DNA Purification Kit (Madison, WI, USA). We amplified four molecular markers through polymerase chain reaction (PCR) using the following primers and annealing temperatures: (1) Cytochrome oxidase I, subunit a (COI): LCO1490 (5’-GGT-CAA-CAA-ATC-ATA-AAG-ATA-TTG-G-3’) and either HCO2198 (5’-TAA-ACT-TCA-GGG-TGA-CCA-AAA-AAT-CA-3’) (Folmer et al. 1994) or C1-N-2776 (5’-GGA-TAA-TCA-GAA-TAT-CGT-CGA-GG-3’) (Hedin & Maddison 2001), 35 × 48–52 °C. (2) Internal transcribed spacer 2 (ITS): ITS4 (5’-TCC-TCC-GCT-TAT-TGA-TAT-GC-3’) and ITS 5.8 (5’-TCG-ATG-AAG-AAC-GCA-GC-3’) (White et al. 1990, Hedin 1997), 35 × 51 °C. (3) Histone 3, subunit a (H3): H3aF (5’-ATG-GCT-CGT-ACC-AAG-CAG-ACV-GC-3’) and H3aR (5’-ATA-TCC-TTR-GGC-ATR-ATR-GTG-AC-3’) (Colgan et al. 1998), 35 × 52 °C. (4) 28S ribosomal RNA (28S): 28S-O (5’-GAA-ACT-GCT-CAA-AGG-TAA-ACG-G-3’) and 28S-C (5’-GGT-TCG-ATT-AGT-CTT-TCG-CC-3’) (Hedin & Maddison 2001), 35 × 50 °C. These markers were complemented by sequences of 16S ribosomal DNA available in GenBank from the study of McHugh et al. (2014). We thus sampled two mitochondrial and three nuclear markers.

We checked the success of PCRs by running the product in 1% agarose gel stained with Biotium GelRed^®^ (Fremont, CA, USA). We purified fragments using a mix of exonuclease I and alkaline phosphatase or washes in NaCl+20% 8000 polyethyleneglycol solution. We sequenced the PCR products in both directions in an ABI 3130xl Genetic Analyzer (Applied Biosystems, Waltham, MA, USA). Chromatograms were individually checked and had poorly sequenced flanks trimmed.

### Sequence alignment and phylogenetic analyses

We aligned sequences of each marker using MUSCLE (Edgar 2004) in MEGA7 (Kumar et al. 2016). We double-checked the alignment of protein-coding genes (COI and H3) for indels and stop codons. The ribosomal markers displayed gappy alignments due to several indels. We alternatively aligned these in the MAFFT web server (Katoh et al. 2018) using the Q-INS-I option and removed poorly aligned regions using the Gblocks 0.9 web server (http://molevol.cmima.csic.es/castresana/Gblocks_server.html) using the “allow gap positions within the final blocks”, “allow less strict flanking positions”, and “do not allow many contiguous non-conserved positions” options. We tested the fit of different evolutionary models (under the Bayesian information criterion) and performed composition homogeneity tests for each partition using IQ-Tree 1.6.12 (Minh et al. 2020). Because many COI sequences failed the homogeneity test, we re-partitioned this marker as codon positions 1+2 and codon position 3 and re-ran the tests. Sequences of the third codon position still failed homogeneity tests, thus we RY-coded this partition for downstream analyses.

We prepared four different matrices: (1) morphological data only (13 outgroups + 119 *Micrathena*), (2) sequence data of the five molecular markers, aligned with MUSCLE, (3) sequence data of the five molecular markers aligned with MAFFT and filtered with gblocks and (4) a total-evidence matrix joining (1) and (2). The sequence-only matrices (2 and 3) include multiple individuals per species, when available, resulting in 20 outgroups (19 spp.) + 221 *Micrathena* terminals (79 spp.). For the total evidence matrix, each species was represented by a single terminal, usually the specimen for which more markers were successfully sequenced. In 11 cases, terminals for a species were prepared by joining sequences from different individuals to reduce missing data; these chimeric terminals were carefully prepared using only individuals shown to be co-specific using matrices 2 and 3, and can be readily identified in Fig. 2 by the presence of two voucher numbers. Additionally, the scorings for *Pronous* combine sequences of *Pronous peje* with morphological scorings of *Pronous tuberculifer*. The total evidence matrix includes 6 outgroups, 79 species of *Micrathena* with both sequence and morphological data and 40 *Micrathena* species which lack sequence data. To check if species lacking sequences were acting as rogue taxa, we summarized the trees of the total-evidence Bayesian analysis twice: one including all taxa, and another excluding non-sequenced taxa using the *exclude* command in MrBayes. A graphical summary of the matrices and phylogenetic methods is available as Supplementary Figure S1.

Each of these four matrices was analyzed under maximum likelihood in IQ-Tree 1.6.12 and under Bayesian inference in MrBayes 3.2.7 (Ronquist et al. 2012). We re-scored polymorphisms as missing entries in the morphological matrix for the maximum likelihood analyses. We ran IQ-Tree using 100 standard bootstrap pseudoreplicates, and the following best-fitting models selected using corrected Akaike information criterion (AIC): MK+G4+F+ASC (morphology), GTR+F+R4 (COI codons 1 and 2), K2P+I+G4 (COI codon 3), K2P+R3 (H3), SYM+R5 (ITS), GTR+F+R4 (28S) and GTR+F+I+G4 (16S). We ran MrBayes remotely in the CIPRES portal (Miller et al. 2010). For each matrix, we performed two independent runs of four chains each, run for 5–10,000,000 generations. We used these models: Mkv + Γ (morphology), GTR + I+ Γ (COI codons 1 and 2, 16S, 28S), SYM + I + Γ (ITS), K2P + I + Γ (COI codon 3), and K2P + Γ (H3). We discarded the first 25% of the samples as burn-in and checked convergence and stationarity with Tracer 1.7 (Rambaut et al. 2018).

### Divergence time estimation

We estimated the age of splits within *Micrathena* by analyzing the total-evidence matrix using a molecular clock approach in BEAST 2.6 (Bouckaert et al. 2019). The few polymorphic scorings in the morphological matrix were re-coded to missing. We ran preliminary analyses using the same models used in the MrBayes analysis, but these failed to obtain adequate sampling for some nuisance parameters of the 16S and third position of COI partitions, even if re-coded as RY; we thus employed a HKY + Γ model for the 16S partition and removed the partition containing the COI third codon position. We used a birth-death tree prior and a lognormal relaxed clock. We employed two calibration points based on the results of Magalhaes et al. (2020). The first is a secondary calibration based on the split between *Micrathena* and *Verrucosa*, estimated to have a 95% interval between 58.8 and 13.6 (mean 34.77) Mya. Thus, we set the age of the node containing *Verrucosa, Pronous* and *Micrathena* with a normal prior with a mean of 37.11 and a standard deviation of 11 (resulting in a 13.2–56.3 Mya 95% interval). The second calibration is based on the age of *Palaeonephila dilitans* Wunderlich, a *bona fide* nephiline preserved in Baltic amber with an estimated age between 47.8 and 43 Myr (see Magalhaes et al. 2020). Nephilidae is closer to Araneidae than to Phonognathidae (see Kallal et al. 2021 and Kuntner et al. 2023), thus we used the age of this fossil to calibrate the root of the tree. We set an exponential prior distribution for this node with an offset at 43 Myr; the mean of the distribution was sampled from a uniform hyperprior with bounds between 0.1 (implying the split between Araneidae and Nephilidae happened immediately before *Palaeonephila*) and 37.6 (implying that only 5% of the prior distribution of the age of the root is older than 156 Myr, the maximum age of Araneidae + Theridiosomatidae found by Magalhaes et al. 2020). We ran two independent chains for 25 million generations each and checked convergence and appropriate sampling of the posterior estimates using Tracer 1.7 after discarding the first 10% of samples as burn-in. The post-burn-in trees of the two chains were combined into a single maximum clade credibility tree.

### Estimation of ancestral ranges and biogeographic events

We modeled the biogeographic history of *Micrathena* using an event-based approach based on the dispersal-extinction-cladogenesis model (DEC; Ree & Smith 2008). We divided the Neotropical region into six areas: Amazonia (M), Andean cloud forests and the Chocó (E), Greater Antilles (I), Atlantic Forest (L), Central and North America (C), and Dry Diagonal (D) (Fig. 1). The limits of each area are adapted from high-resolution shapefiles by Olson et al. (2001) and Romano (2017). Andean cloud forests and Chocó were considered a single area because there is only a single known species endemic to lowland Chocó (*M. guayas*). We then classified each species of *Micrathena* as either present or absent in each area. A database of species distributions was made from records from taxonomic publications on the genus (Levi 1985; Gonzaga & Santos 2004; Magalhaes & Santos 2011, 2012; Magalhaes et al. 2017) and from specimens examined by us from recent collections. We also included iNaturalist (https://www.inaturalist.org/) records checked by us (up to 29/07/2020) and whose identity could be unambiguously determined from photos. In only two species there are relevant range extensions when compared to published work: we have examined specimens of *M. furcula* from the Amazon and Atlantic Forests, and of *M. patruelis* from the Amazon Forest, and thus these species have been scored as present in these areas. We pruned outgroups from the biogeographic analyses because both *Pronous* and *Verrucosa* are diverse genera whose species occur in different parts of the Neotropics and there is no species-level phylogeny of them; thus, including the few species of these genera sampled in our matrix could bias the ancestral range estimated for *Micrathena*.

We estimated ancestral ranges in the R package BIOGEOBEARS (Matzke 2013). First, we explored the fit of the data to six alternative biogeographic models (DEC, DIVA-like and BayArea-like, and their variants including a J founder-event speciation parameter). Each of these six models were run with an unconstrained dispersal matrix (dispersal multipliers among all areas set to 1) and with a dispersal matrix that takes distances between areas into account (dispersal multipliers between: adjacent areas, 1; areas separated by another area or the sea, 0.5; areas separated by two areas or one area and the sea, 0.1). In the latter case, we included a free “w” parameter to estimate the weight of dispersal multipliers.

To test whether closure of the Isthmus of Panama played a role in *Micrathena* biogeography, we re-run the favored model under a time-stratified model in three different scenarios: (1) no South American areas before the formation of the Isthmus (forcing all *Micrathena* to occur in Central and North America before that), (2) no Central and North America before the formation of the Isthmus (forcing all *Micrathena* to occur in South America before that), and (3) reduced dispersal between Central and South America before formation of the Isthmus (all dispersal multipliers between these areas reduced to a tenth of current values). We used a very conservative age estimate of formation of the Isthmus of 11 Mya since there are significant bouts of faunal exchange between these landmasses as early as 8 Mya (O’Dea et al. 2016).

We tested all the models outlined above using the maximum clade credibility tree estimated with BEAST. However, both the topology and nodal ages were estimated with considerable uncertainty, especially because of those taxa lacking sequence data. Thus, we also estimated ancestral ranges over a sample of 100 trees from the posterior distribution of the BEAST analysis using the script by Magalhaes et al. (2021). We then estimated the biogeographic transitions and their timings using 10 biogeographic stochastic mappings (Dupin et al. 2017) for each of the 100 trees, resulting in 1,000 different biogeographic histories. We calculated the average number of biogeographic events (e.g., dispersals among areas, speciation) in each of the histories, discriminated by area. We discriminate three different types of speciation: *in-situ speciation* refers to when an ancestor inhabiting a single area splits into two descendants inhabiting the same area; *allopatric speciation* refers to events where a widespread ancestor inhabiting two or more areas splits into descendants occupying mutually exclusive smaller areas; *subset sympatry speciation* refers to events where an ancestor inhabiting two or more areas splits into one descendant with identical distribution to the ancestor and another inhabiting a single area. Finally, we calculated how the rates of dispersal and in-situ speciation changed through time; for this, for each replicate we counted the number of such events in 1-million-years bins and corrected this value by the number of *Micrathena* lineages at the beginning of each bin. We also estimated the total time each area was inhabited by summing the length of the branches of the lineages occupying it. We report average values of each of the 1,000 biogeographic histories.

Our dataset includes all *Micrathena* species, 40 of which lack DNA sequences and whose phylogenetic position was inferred from morphology alone. In most recent biogeographic studies, species lacking sequences are not included in the sampling and thus any results ignore the effect they might have on the inference. To simulate this situation, we re-ran the ancestral range estimation and biogeographic stochastic mapping after removing these 40 species by pruning them from the trees and compared the results with those of the full dataset.

### Diversification in lowlands and highlands

The previous analyses indicate that in-situ speciation is especially high in the Andean cloud forests (see Results below). Thus, we tested whether clades occupying highlands have higher diversification rates than those in lowlands using the geographic state-dependent speciation and extinction model with hidden states (GeoHiSSE; Caetano et al. 2018). We classified *Micrathena* as occurring in highlands, lowlands, or both by extracting elevations for each occurrence record in our database from an elevation raster using the sf and terra packages for R (Pebesma & Bivand 2023). Species were classified as occupying highlands if at least 25% of their records occur in areas 1200 meters above sea level (following Pérez-Escobar et al. 2022 as the minimum elevation for cloud forests), and as occupying lowlands if at least 25% of their records occur below that threshold; six species fulfilled both criteria and were classified as widespread. Net diversification in highlands and lowlands was estimated using seven different models in the *hisse* package for R (Beaulieu & O’Meara 2016): (1) diversification rates equal among highlands and lowlands, no hidden area, (2) same, but with a hidden area, (3) same, but with four hidden areas, (4) diversification rates allowed to be different among lowlands and highlands, no hidden area, (5), same, but with a hidden area, (6) same, but with four hidden areas and (7) same as 2, but with no cladogenetic events allowed (MuSSE-like model). These seven alternative models allow to test if net diversification differs among lowlands and highlands while also allowing for diversification being explained by other factors (“hidden” areas); we used a maximum of four hidden areas as this matches the same number of areas of the BIOGEOBEARS analysis. To account for phylogenetic uncertainty, we estimated diversification parameters for each of these models in 11 different trees (the MCCT, and 10 trees randomly drawn from the posterior distribution of the BEAST analysis). We averaged the estimated diversification rates in each area across all models and trees, weighting by the AIC weight of each model.

## Results

### Phylogeny of *Micrathena*

All analyses including DNA sequences result in *Pronous* and *Verrucosa* as the two successive sister-groups of *Micrathena*; morphology alone produces ambiguous results regarding the closest relatives of the genus. Our analyses include sequences of the Neotropical genera *Aspidolasius, Xylethrus, Spinepeira* and *Encyosaccus* for the first time in the literature; none of these are recovered in the micrathenine clade (sensu Scharff et al. 2020) (Suplementary Figures S3–4, 7–8). All data matrices support *Micrathena* monophyly (Fig. 2, Supplementary Figures S2–S9). They also indicate that the bulk of *Micrathena* is divided into two lineages, one containing the *swainsoni* clade (*swainsoni, triangularispinosa* and *militaris* species groups), and the other unites the remaining species (the *gracilis*, *clypeata* and *saccata* clades, each of these also well-supported across analyses) (Fig. 2). *Micrathena spitzi* is either recovered as the sister to all other species in the genus (Fig. 2) or as sister to the *schreibersi* + *cornuta* groups (Fig. 3).

**Figure 3.**
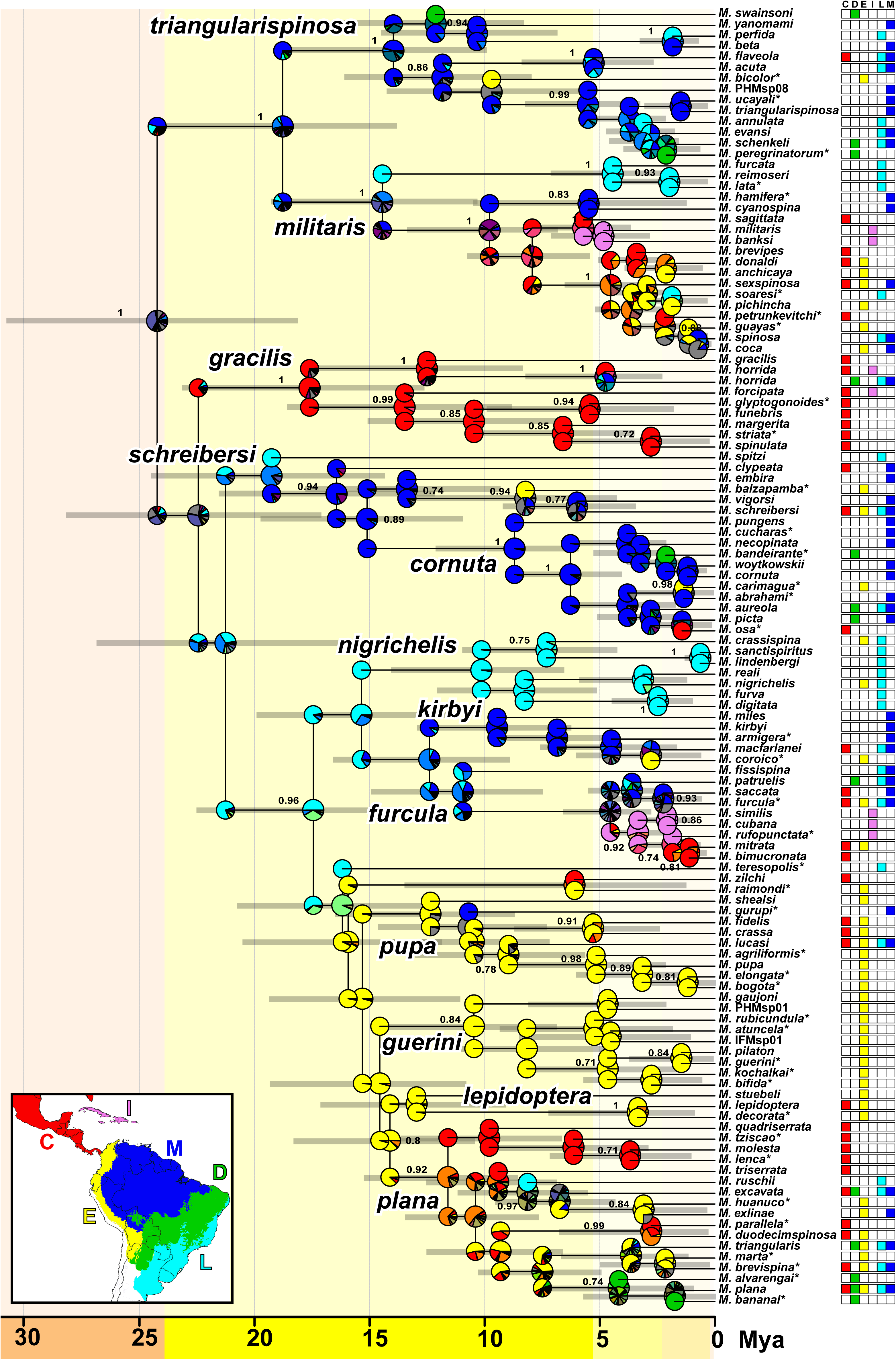
Dated maximum credibility tree estimated using Bayesian inference and including all species of *Micrathena*. Species marked with an asterisk only have morphological data available. Values on branches are posterior probabilities; only values larger than 0.7 are shown. Grey bars are 95% highest posterior density intervals of node ages. Pies at nodes are probabilities of ancestral ranges estimated under the dispersal-extinction-cladogenesis model. Area codes: C = Central and North America, E = Andean forests and Chocó, D = Dry Diagonal, I = Greater Antilles, L = Atlantic Forest, M = Amazon.

The datasets containing only morphological data support monophyly of most of the previously recognized *Micrathena* species groups, except for the *triangularispinosa*, *furcula* and *guerini* groups. These results are mirrored by molecular data. However, the relationships among species groups differ between morphology-only (Supplementary Figures S2, S6) and the other datasets, and only the latter will be discussed in detail. Analyses of the sequence-only datasets produced nearly identical results among them, regardless of alignment algorithm, removal of poorly aligned regions, or method of phylogenetic inference (Supplementary Figures S3, S4, S7, S8). These results closely mirror those of the total-evidence datasets (Fig. 2, Supplementary Figures S5, S9). The few conflicting clades are listed below.

Phylogenies of all taxa (including non-sequenced *Micrathena* species) result in poorer nodal supports of some of the shallower clades; the 40 taxa lacking molecular data act as rogue taxa and erode the supports of otherwise robust nodes. Re-summarizing the trees of the total-evidence dataset, but excluding non-sequenced species, substantially increases posterior probabilities across the tree (compare Fig. 2 and Supplementary Figure S9, which are based on the exact same analysis). However, it is noteworthy that the four main clades and several of the *Micrathena* species groups are well-supported even upon inclusion of those species.

Several *Micrathena* species groups as re-delimited by Magalhães & Santos (2012) are monophyletic, including the *militaris*, *plana*, *kirbyi*, *cornuta*, and *lepidoptera* species groups. The *triangularispinosa* and *gracilis* groups are paraphyletic because the monotypic *swainsoni* and *funebris* groups nest within them, respectively; it should be noted that the *triangularispinosa* group includes *M. beta* and *M. perfida*, two species with male genital morphology resembling that of the *militaris* group. The *furcula* group is diphyletic; a well-supported clade reunites most of its species, but *M. digitata* and *M. furva* are more closely related to *M. nigrichelis* and other species endemic to the Atlantic rainforest. The *cornuta* group is nested within the *schreibersi* group. The monophyly of the latter is further challenged by the unstable position of *M. spitzi*: this morphologically aberrant species was recovered in most datasets as sister to all other *Micrathena* with good support. Morphology-only datasets support it as part of the *schreibersi* group, and trees from the dated analysis show ambiguous signal: it is recovered as part of the *schreibersi* group in ∼43% of the posterior trees, and in the basal split within the genus in the remainder. Finally, the *guerini* species group is polyphyletic and intertwined with many species previously considered as *incertae sedis*. Most of these species are now grouped in at least three major clades (labeled *nigrichelis*, *pupa* and *guerini* in Figs 2, 3), but some species remain orphan lineages without close relatives (e.g., *Micrathena zilchi*, *M. stuebeli*). The *nigrichelis* clade contains species mostly from the Atlantic Forest and it is a close relative of the *kirbyi* and *furcula* species groups. However, this clade is only recovered in datasets aligned with Muscle (Fig. 2, Supplementary Figures S3, S7). The *pupa* and *guerini* clades contain mainly Andean endemics and are closely related to the *lepidoptera* and *plana* species groups.

### Species limits in *Micrathena*

While not a major aim of this study, we observed that morphology-based species identifications are almost ubiquitously supported by sequence data (either using COI, ITS or the multi-locus datasets). Most species are monophyletic (Supplementary Figures S3– S4, S7– S8, S10). The few exceptions are the pairs *M. mitrata–M. bimucronata* and *M. triangularis–M. plana*, which indicates that these species may need their taxonomy revised, and the pair *M. sanctispiritus*–*M. lindenbergi*, whose extremely shallow divergence suggests these two may be synonyms. Conversely, at least two species display deep intraspecific divergences that indicate potential cryptic diversity: *M. lucasi* and *M. horrida*. In the latter case, one of the clades is exclusively South American, while the other occurs in Central America and the Antilles. For this reason, these two clades of *Micrathena horrida* were considered as independent lineages for downstream biogeographic analyses; although there are many available names currently synonymized under *M. horrida* that may be used for these clades (e.g., *Acrosoma longicauda* O. Pickard-Cambridge, 1890), diagnosing them morphologically is difficult and we refer to both clades as *M. horrida* until the relevant types are examined and the species is properly re-delimited.

### Divergence time estimation

*Micrathena* split from its sister genus *Pronous* between 40– 25 Mya and began diversifying between 30–20 Mya (Fig. 3, Supplementary Figure S11). Estimates of pairwise divergence rates of H3 (0.33% / Myr) and 16S (1.23%) are within the same order of magnitude as those estimated for other spiders (see Bidegaray-Batista and Arnedo, 2011), while that of 28S (1.50%) is ∼12 times faster, although the rates may not be comparable since the primers used to amplify this gene are not the same in both studies. COI divergence rates are not reported because this gene was partitioned by codon.

### Estimation of ancestral ranges and biogeographic events

Results of the ancestral range estimation are qualitatively similar among different biogeographic models, except for the BayArea-like models. The best-fitting model is that using the DEC+J model including a dispersal matrix that takes distances among areas into account. The DEC+J and DEC models yield qualitatively identical results, except for a few nodes where DEC+J estimates a founder-event speciation, and DEC estimates anagenetic dispersal followed by allopatry. Thus, they differ only in the dispersal event happening at a node or along the branch immediately before it. We report results of the DEC model since (1) it is more straightforward to summarize dispersal events in this model (since it only includes anagenetic dispersal, as opposed to the cladogenetic and anagenetic dispersal in DEC+J), and (2) because the age of biogeographic events in models with the J parameter are more strongly tied to the ages of nodes (Magalhaes et al. 2021). Ancestral range estimation indicates that several clades of *Micrathena* have diversified in a single area (Fig. 3): e.g., the *schreibersi* + *cornuta* groups are mainly from the Amazon, the *gracilis* group mainly diversified in Central America and the *pupa* and *guerini* groups in the Andean forests. The ancestral range of the whole genus was most likely wide, consisting of M+L+C or M+L+E+C (see Fig. 1 for area codes).

The areas with more species of *Micrathena* are the Andean forests and Chocó (43 spp.) and the Amazon (42), followed by Central America (37), the Atlantic Forest (33), the Dry Diagonal (13), and the Antilles (7) (Table 1). The Andean forests have the most in-situ speciation events (21.6 events in average across the 1,000 possible biogeographic histories); despite the similar absolute richness, in situ-speciation in Amazon is 20% lower (17.4) than in the Andes (Fig. 1). However, the Amazon shows higher overall speciation, and is also the major source of species to other areas; it sourced ∼57% more species (18.7) than the Andes (11.9) (Table 1, Fig. 1). The Dry Diagonal has the lowest in-situ speciation (0.38), indicating that its species either arrived from other areas (10.6) or speciated from widespread ancestors allopatrically (3.8) or via subset sympatry (2.0). The Antilles have fewer species than the Dry Diagonal, but higher in-situ speciation (2.6), which is consistent with the observation that at least two clades (the *furcula* and *militaris* groups) radiated after reaching the islands. Generally, dispersers incoming from other areas contribute more to the diversity of each area than does in-situ speciation (Table 1), with the exception of the Amazon and Andean forests. Allopatric speciation most frequently happened between Central America and Andean forests (9.6 events) or between the Amazon and other areas (Atlantic Forest, 8.6; Andean forests, 8.4; Dry Diagonal, 5.33; Central America, 4.04).

*Micrathena* have dispersed to the Greater Antilles 3–5 times independently, most likely from Central and North America or larger ranges containing this area (76% of the events), and these events most likely took place in the last 20 Myr, thus much after the hypothesized age for the existence of GAARlandia (∼33 Mya) (Fig. 4). This inference is robust to phylogenetic and age uncertainty. Faunal interchange between South and Central America was frequent and started as early as 30 Mya; rates are asymmetric, with dispersals from South to Central America more common (10–25 events) than the converse (2–15) (Fig. 4). The majority (63%) of dispersal events between the subcontinents predates formation of the Isthmus of Panama (∼3.5 Mya), with only 37% of such events happening after 3 Mya. Biogeographic models that preclude (or hamper) dispersal before formation of the isthmus fit the data much worse than an unconstrained model (Supplementary Figures S12–S14) and can be rejected (AIC weight of the constrained models lower than 0.001 in all cases). Finally, rates of dispersal have not increased substantially after formation of the isthmus (Supplementary Figure S15). Interchange between the Amazon and Atlantic forests, or between the Atlantic Forest and the Andean forests, has also happened repeatedly throughout the history of *Micrathena* (Fig. 4), although in this case rates have also been asymmetric: we infer ∼5 dispersal events from the Amazon to the Atlantic Forest and ∼2 in the reverse direction, and ∼2 dispersal events from the Atlantic Forest to the Andes and ∼1 in the reverse direction.

**Figure 4.**
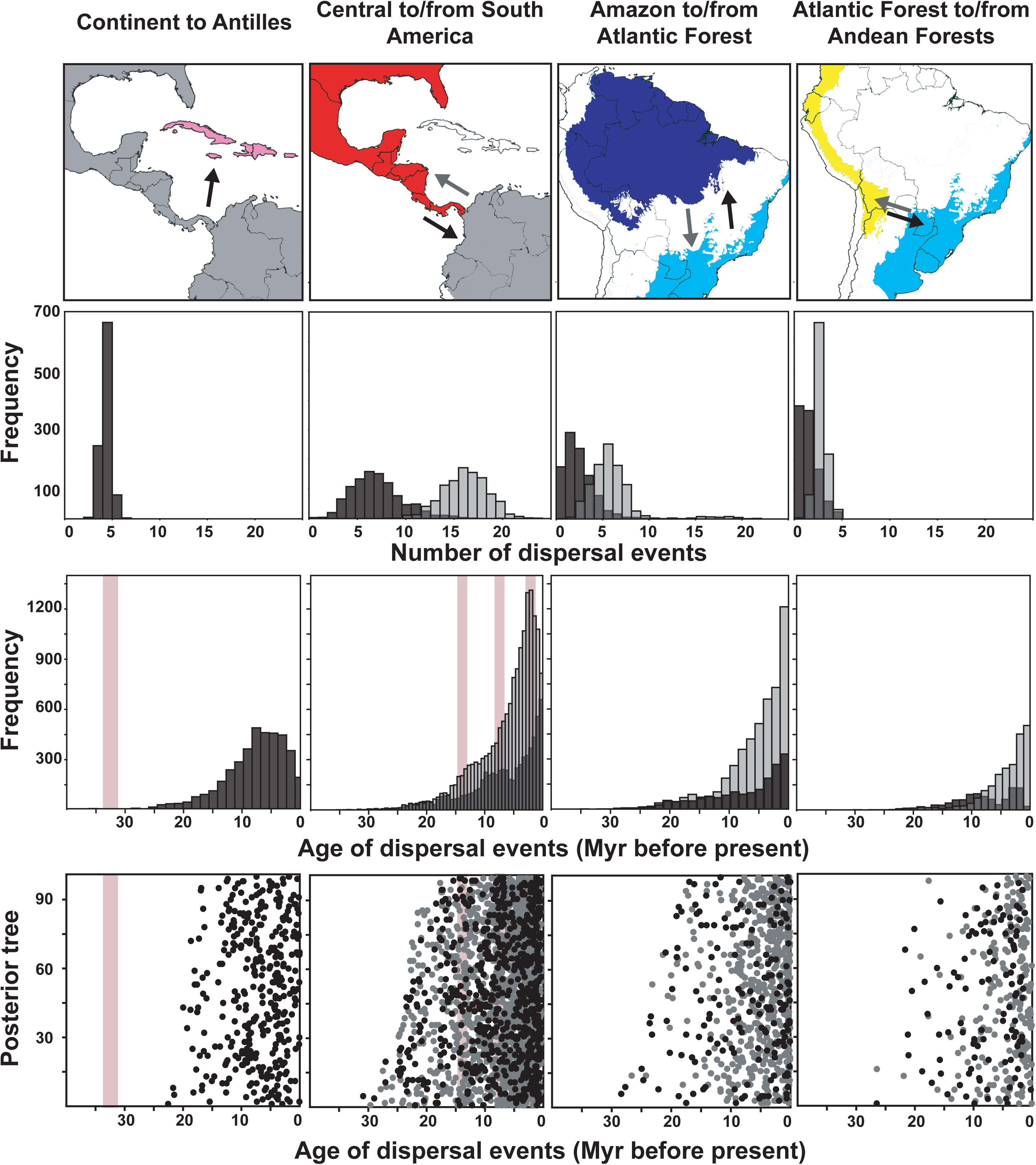
Age and frequency of dispersal events of *Micrathena* between selected areas in the Neotropical region: first column, from the continent to the Antilles; second column, from Central to South America (black) and vice-versa (gray); third column, from the Atlantic Forest to the Amazon (black) and vice-versa (gray); fourth column, from the Andean forests to the Atlantic Forest (black) and vice-versa (gray). Estimates are based on 10 biogeographic stochastic maps for each of 100 trees from the posterior distribution of the dated analysis. Second row: frequency of dispersal events across the 1000 stochastic maps. Third row: frequency of the age of dispersal events across the 1000 stochastic maps. Fourth row: each dot represents a dispersal event; each unit in the Y axis represents a stochastic map of a different tree from the posterior distribution. Pink bars represent the period of existence of GAARlandia in the first column, and the multiple hypotheses of timing of closure of the Panama Isthmus in the second column.

The biogeographic results described above refer to the analyses sampling all *Micrathena*, including those lacking sequences. We compared these with the results obtained by pruning the 40 species which lack DNA sequences. When non-sequenced species are excluded, in-situ speciation in the Amazon (14.4) is inferred to be 36% higher than in Andean cloud forests (10.6) (Table 1), which contradicts the results obtained with the full dataset. This may be explained in part by geographic bias in the sampling: we could sequence more than 75% of the species present in all areas, except for the Andes, where only 55% of the species were sampled. This impacts the conclusions drawn from the two datasets. Both indicate that species richness in the Andes started rising sharply around ∼17 Mya. However, the full dataset indicates this area became the most species-rich at ∼5 Mya, while in the dataset containing only sequenced species the Andes end as only the fourth most diverse area (Fig. 5). The full dataset indicates that in-situ speciation rates in the Andes increased above average twice, around ∼17 and 6 Mya, surpassing in-situ speciation rates at the Amazon (Fig. 5). However, this pattern is not observed in the pruned dataset: a single increase in in-situ speciation in the Andes is detected at 17 Mya, and it never surpasses the rates of the Amazon (Fig. 5).

**Figure 5.**
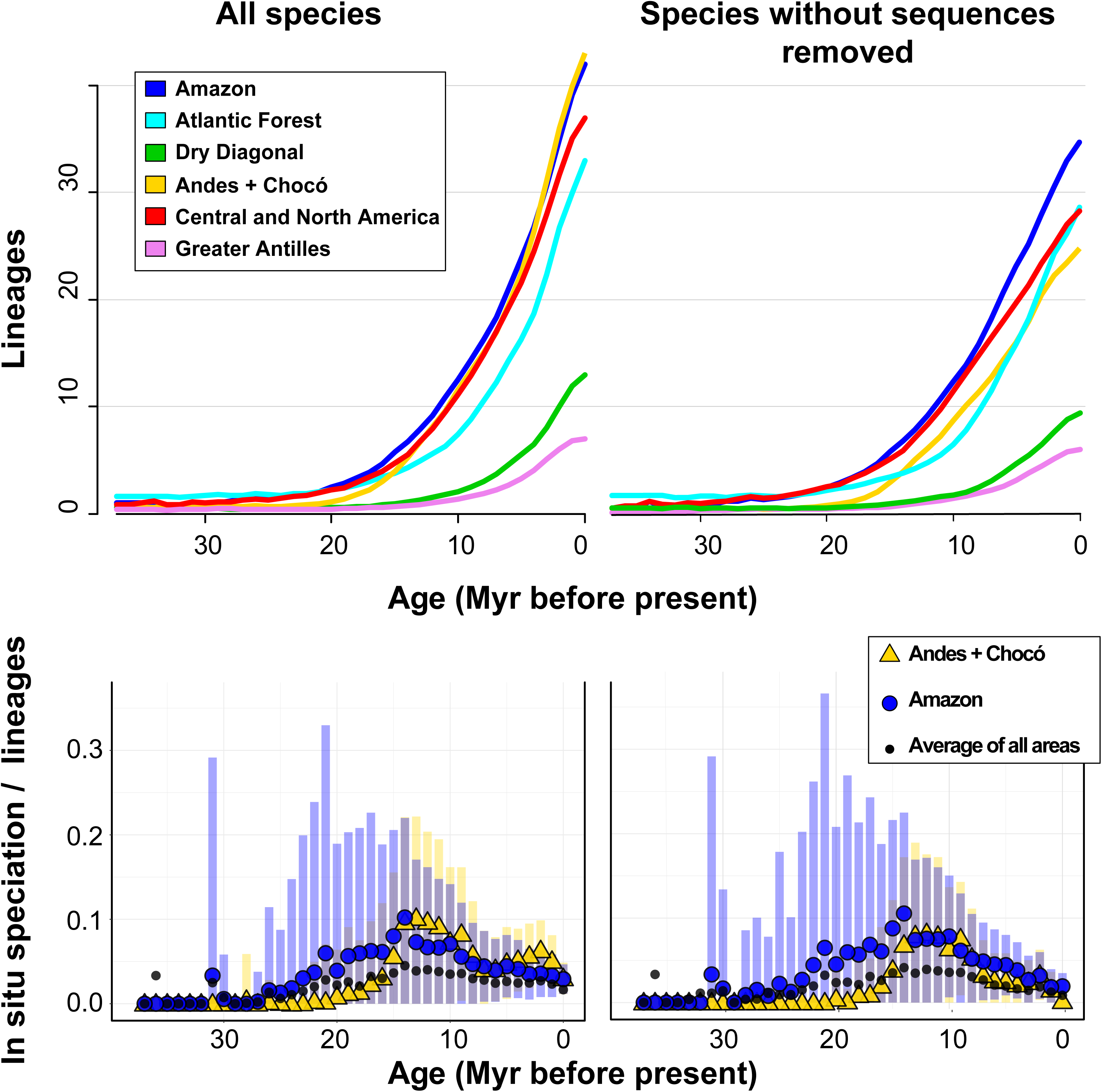
Failing to sample all species may hamper our understanding of diversification patterns. Top row: number of species of *Micrathena* in each area through time. Only when including all species (left) we are able to see that the Andean forests and Chocó are the most diverse areas; including only sequenced species (right) leads us to think that the Amazon is the most diverse area. Bottom row: diversification rates of *Micrathena* in the Amazon (blue) and in the Andean and Chocó forests (yellow); dots are average values and bars are standard deviations. Only by including all species (left) it was possible to observe two bouts of diversification in the Andes and Chocó, around ∼17 and ∼4 million years ago; including only sequenced species (right) leads us to think that the Amazon always had higher diversification rates than the Andes and Chocó.

### Diversification in lowlands and highlands

Among the eleven trees used for estimating net diversification with GeoHiSSE, eight favored models 1 or 3 (where net diversification is equal in lowlands and highlands), and three favored model five (where net diversification may be different among these areas); even in the latter case, net diversification of highlands is between 77.7–82.3% of that in lowlands, and thus within the same order of magnitude. After averaging the two parameters across all models and trees, and weighting by the AIC weight of each model, we found that net diversification in the highlands is 0.097 while in lowlands it is 0.107, and thus nearly the same. Thus, clades that shifted to highlands maintained similar net diversification rates as those that stayed in lowlands.

## Discussion

### *Micrathena* relationships

Coming from a long way that started 40 years ago in the hand-drawn cladogram by Levi (1985), we reached a new phylogenetic hypothesis for *Micrathena* based on morphological and sequence data and including all its known species. Considering the high diversity and geographical extension of the genus, and the paucity of complete or near-complete invertebrate phylogenies in the Neotropical region, our work has few parallels in the literature (e.g., Cameron et al. 2007, Caterino & Tishechkin 2015, Ojanguren-Affilastro et al. 2016, Price et al. 2016, Cabra-García & Hormiga 2020). Morphology and sequence data are generally congruent, and the main clades and species groups are well-supported. This suggests that the placement of non-sequenced species is reliable; the topological instability that leads to poor nodal support is mainly restricted to small groups of closely related species. We recover *Pronous* as the sister group of *Micrathena*, as previously suggested by Cabra-García & Hormiga (2020). This genus can be regarded as part of the micrathenines *sensu* Scharff et al. (2020). Morphological synapomorphies uniting the two genera are the rectangular abdomen and three pairs of lateral abdominal spines. The new phylogenetic hypothesis calls for a taxonomic rearrangement of species groups within *Micrathena* as they are currently recognized; some of them are paraphyletic, and the *guerini* group needs to be radically re-delimited, as most of its species now belong in three different clades. The main open question regarding *Micrathena* phylogeny is the position of *M. spitzi*. Our results inconsistently place this species either as part of the *schreibersi* group, where it was originally placed by Levi (1985), or as sister to all other *Micrathena*. If confirmed, the latter would call for re-interpretations of the evolution of morphology within the genus.

### Sources and sinks of *Micrathena* diversity

*Micrathena* diversification took place relatively recently and rapidly, as its ∼120 species started diversifying between 30 and 20 Mya. We found the Amazon to be the area where most species of the genus originate, but only when the three speciation processes are considered jointly: in-situ speciation, allopatric speciation, and subset-sympatry speciation. The Amazon is also the main source of species of *Micrathena* to other areas through dispersal, a common pattern in most taxonomic groups surveyed (Antonelli et al. 2018b). These results may be partly explained by its large area and extensive borders with other regions, by its historical climatic stability in comparison with other Neotropical forests (Costa et al. 2017), and also by the long-term occupation of this area (Antonelli et al. 2018b); in *Micrathena*, the Amazon is also the area occupied for a longer period.

On the other extreme, the Dry Diagonal has the lowest in-situ speciation. It does have some endemic species (e.g., *M. swainsoni, M. peregrinatorum, M. bandeirante, M. bananal*) that apparently originated via allopatric or subset-sympatry speciation, with their sister groups inhabiting the Amazon. This scenario is consistent with cycles of expansion and retraction of humid forests over the Dry Diagonal (e.g., Auler et al. 2004). The Dry Diagonal also sources very little species to other areas, despite its large area of contact with neighboring regions. Thus, the Dry Diagonal acts as a sink of *Micrathena* diversity. Consistent with this, shifts from mesic to xeric habitats are more frequent than the converse in many groups (e.g., Antonelli et al. 2018b, Ziska et al. 2020, Buianain et al. 2022). Finally, perhaps some of the species recorded as present in the Dry Diagonal may be accidental records taken from gallery forests or *brejos de altitude* (e.g., *M. excavata, M. aureola, M. horrida*); this type of mesic vegetation that borders water bodies has long been recognized as an important corridor of biota that is otherwise restricted to rainforests (e.g., see Costa 2003).

Mountain uplift may lead to high species richness through increases in speciation following orogenesis (e.g., Xing & Ree 2017). Our results show that the Andean forests have the highest number of *Micrathena* species and the highest levels of in-situ speciation, despite its smaller area in comparison with the Amazon. Our results indicate at least two bouts of speciation in the Andes, at ∼17 and ∼5 Mya (Fig. 5). These ages are consistent with the timing their uplift: by the Miocene the mountain range was 1500–2000 m high (Hartley 2003, Graham 2009), and orogenesis has been particularly rapid and recent in the northern Andes (Pérez-Escobar et al. 2022), where most of Andean *Micrathena* occurs. This allowed the establishment of cloud forests with different environmental conditions from the surrounding lowland forest and the onset of montane taxa diversification in the region between 20–10 Mya (Hoorn et al. 2010). It also resulted in the appearance of geographical barriers that limit dispersal and are coincident with areas of endemism (Hazzi et al. 2018). Thus, we interpret that Andean uplift promoted *Micrathena* speciation. The GeoHiSSE results indicate that net diversification is similar between lowland and highland species, and thus it seems that orogeny has not accelerated diversification rates in this genus. However, the high in-situ speciation of Andean *Micrathena* may be taken as evidence that this region generates a high number of endemic species. These may be (1) narrow-ranged and topographically trapped in small, isolate mountain tops or valleys (e.g., *M. pichincha, M. kochalkai, M. bicolor*), or (2) ecologically adapted to the particular climatic conditions of cloud forests, occurring in a wide range but always in mid to high elevations (e.g. *M. crassa, M. agriliformis, M. lepidoptera*). However, the mechanisms by which orogenesis lead to an increase in speciation are not fully understood (Vargas et al. 2020).

A complete taxonomic sampling was paramount to correctly identify the main centers of speciation in *Micrathena*. When non-sequenced species are excluded, the exceptional in-situ speciation in the Andes is masked (Fig. 5), precluding the link between orogeny and diversification of this group. This highlights the importance of striving to produce complete taxonomic samplings, especially when trying to understand diversification dynamics. We hypothesize that precisely those areas that generate the most diversity will harbor a large quantity of rare and short-ranged species that are difficult to find and sample for DNA extraction. If this is true, many studies based only on molecular data will miss an important proportion of species from these speciation nexuses, which may hamper their identification as such. It is noteworthy that other studies that also achieved complete or near-complete taxonomic sampling have used taxonomy or morphology as proxies to constrain the placement of non-sequenced species (Upham et al. 2019, Landis et al. 2021) or employed total-evidence analyses relying on morphology to place non-sequenced species (Price et al. 2016, Ojanguren-Affilastro et al. 2016, Cabra-García & Hormiga 2020, Matos-Maraví et al. 2021). Both approaches highlight the importance of morphology to estimate the phylogenetic position of rare species whose molecular data is wanting.

### South American isolation: GAARlandia and closure of Panama Isthmus

We estimate that four *Micrathena* lineages reached the Greater Antilles independently: *M. horrida, M. forcipata*, and representatives of the *militaris* and *furcula* groups, corroborating earlier findings that showed multiple dispersal events to the islands (McHugh et al. 2014, Crews & Esposito 2020, Shapiro et al. 2022). Our more comprehensive geographic and taxonomic sampling allows us to identify Central and North America as the most likely source of the lineages reaching the Antilles. The timing of these dispersals is never older than 20 Myr, and thus much younger than the hypothesized age of GAARlandia (33 Myr; see Iturralde-Viñent & MacPhee 1999). Taken together, these results indicate that *Micrathena* reached the Greater Antilles via over-water dispersal from Central or North America, and not via a land bridge connecting the islands to South America.

We also infer over-water dispersal between South and Central America: we estimate 17–31 dispersal events between these continents, and most predate the final closure of the Isthmus, with some as old as 30–15 Mya. Forcing dispersal to happen only after a conservative 11 Mya threshold results in very poor models, mainly because it results in several additional dispersal events. By using the more generally accepted age of 3.5 Myr for the Isthmus of Panama would result in even poorer models. Finally, dispersal rates between South and Central America remained nearly constant over time. Thus, our data clearly indicates that *Micrathena* did not rely on the Panama Isthmus to cross between these two subcontinents. Notably, in many flying insect groups, dispersal between these continents is coincident or has increased with the formation of the isthmus (Hines 2008, Matos-Maraví et al. 2021, Price et al. 2022). *Micrathena* are not particularly good dispersers: we found that several clades speciate in a single geographic area; phylogeographic analysis of insular species indicates that each lineage is usually endemic to a single island (Shapiro et al. 2022); and the genus has not been recorded to disperse through ballooning—Bell et al. (2005) state that *Micrathena sagittata* has been recorded ballooning, but the work they cite as the source of this information never mentions this species (Decae 1987). How they were able to disperse over water, even when different clades of winged insects could not, is intriguing.

### Recurring connections among Neotropical rainforests

Dispersal from the Amazon to the Atlantic Forest was the most important dispersal route we identified; it is especially notable considering they are not adjacent. This reinforces the well-established pattern of historical connection between these two areas (Batalha-Filho et al. 2013, Ledo & Colli 2017, Peres et al. 2017, Prates et al. 2016, de Sá et al. 2019, Matos-Maraví et al. 2021, Nascimento et al. 2021). We infer that 1–5 of these dispersal events happened in the last two million years and may be linked to Pleistocene humid periods that lead to the replacement of dry forests by rainforests in presently xeric areas of northeastern Brazil (Auler et al. 2004). Alternatively, some species use the gallery forests in the Dry Diagonal as a corridor between the two biomes (Costa 2003). These hypotheses may be evaluated through phylogeographic studies of selected species. It should be noted that the number of dispersal events inferred here may be an underestimate: several species occur in both the Amazon and Atlantic Forest (e.g., *M. schreibersi, M. aureola, M. acuta*), and may have a complex phylogeographic history in these biomes. The route connecting the Atlantic Forest to the Andes is not as important, but we noted an interesting pattern: it usually connects species from the southern Atlantic Forest to the southern limit of the Andean cloud forests (e.g., *M. nigrichelis, M. crassispina*). This is also consistent with Pleistocene expansion of forest-associated taxa in the southern end of the Atlantic Forest, as has been described for other taxa (Trujillo-Arias et al. 2017, 2020).

## Conclusions

1. We propose a new and robust phylogenetic hypothesis for *Micrathena*. Morphology and sequence data are generally congruent and allow the recognition of several well-delimited species groups. The ∼120 species of the genus began diversifying around 30–20 million years ago and is most diverse in Neotropical rainforests, particularly the Amazon and Andean cloud forests.
2. We estimated that in-situ speciation is highest in the Andean forests, and lowest in the Dry Diagonal. In-situ speciation in the Andes increased between 17 and 5 million years ago, which is coincident with periods of intense orogeny, suggesting that mountain uplift may have promoted diversification of these spiders. Despite having lower in-situ speciation rates than Andean forests, the Amazon display the highest overall speciation rates and is also the most frequent source of species dispersing to new areas.
3. *Micrathena* reached the Greater Antilles at least four independent times, always from Central and North America. Our estimates of the age of such dispersal events are never older than 20 Mya, indicating that these spiders reached the islands through over-water dispersal, not through the hypothesized Oligocene GAARlandia land bridge.
4. Dispersal events between South and Central America were frequent but asymmetric; we estimate ∼23 of such events, but dispersals from South to Central America were twice as frequent as the converse. Dispersals between these continents started as early as 30–20 million years ago and remained constant through time, indicating that this interchange precedes the closure of the Panama Isthmus, and was unaffected by it.
5. Dispersal between Neotropical rainforests, such as between the Amazon and the Atlantic Forest, or the Atlantic and Andean cloud forests, have been frequent over the last ∼20 million years, although rates have also been asymmetric. Connections between the Atlantic and Andean forests usually occurred through the southern portion of the Andes.
6. Some of our conclusions could only be reached by including all species in the phylogeny; for instance, using only sequenced species failed to reveal the increase of in-situ speciation in the Andes. This highlights the importance of complete phylogenies when testing biogeographic hypotheses, and the usefulness of morphology in estimating the relationships of species for which molecular data is unavailable.

## Acknowledgments

We thank Lorenzo Prendini and Pio Colmenares (AMNH), Lauren Esposito (CAS), O. Francke and A. Mondragón (CNAN-Ar), Antonio Brescovit and C. Rheims (IBSP), M. Ramírez (MACN-Ar), Gonzalo Giribet and Laura Leibensperger (MCZ), Alexandre Bonaldo (MPEG), Diana Silva (MUSM) and Ricardo Pinto da Rocha (MZSP) for loaning specimens from their collections and welcoming us into their institutions. Specimens or sequences of *Micrathena* were kindly collected and made available by G. Barrantes, J. Cabra-García, F. Álvarez Padilla, and F. Labarque. We thank Arthur Anker, Alfredo D. Colón Archilla, Alexis Callejas Segura, César Favacho, Gernot Kunz, and Ricardo Arredondo by the photographs used in Fig. 2. Philip Russo provided literature records of *Micrathena* species. ILFM visits to the AMNH, MCZ and CAS collections was supported by an Ernst Mayr Travel Grant from the MCZ and a Theodore Roosevelt Memorial Grant from the AMNH. Earlier versions of this work were presented in the Reunión Argentina de Cladística y Biogeografía thanks to an invitation by Dolores Casagranda. We thank all the observers who uploaded *Micrathena* observations to iNaturalist, as well as the platform itself for making the data readily available. We thank J. Cabra García for thoughtful comments on an early draft and G.H.F. Azevedo for suggesting the GeoHiSSE analysis. This research was supported by a CONICET postdoctoral fellowship to ILFM and by grants from Fondo para la Investigación Científica y Tecnológica (PICT 2020-1907), Consejo Nacional de Investigaciones Científicas y Técnicas (PUE-098), Conselho Nacional de Desenvolvimento Científico e Tecnológico (407288/2013-9, 405795/2016-5, 311843/2022-0), Fundação de Amparo à Pesquisa do Estado de Minas Gerais, Brazil (PPM-00605-17), Instituto Nacional de Ciência e Tecnologia dos Himenópteros Parasitóides da Região Sudeste Brasileira, Brazil (http://www.hympar.ufscar.br/), and Fundação de Amparo à Pesquisa do Estado de São Paulo, Brazil (2011/50689-0).

---

**Figure S1.**
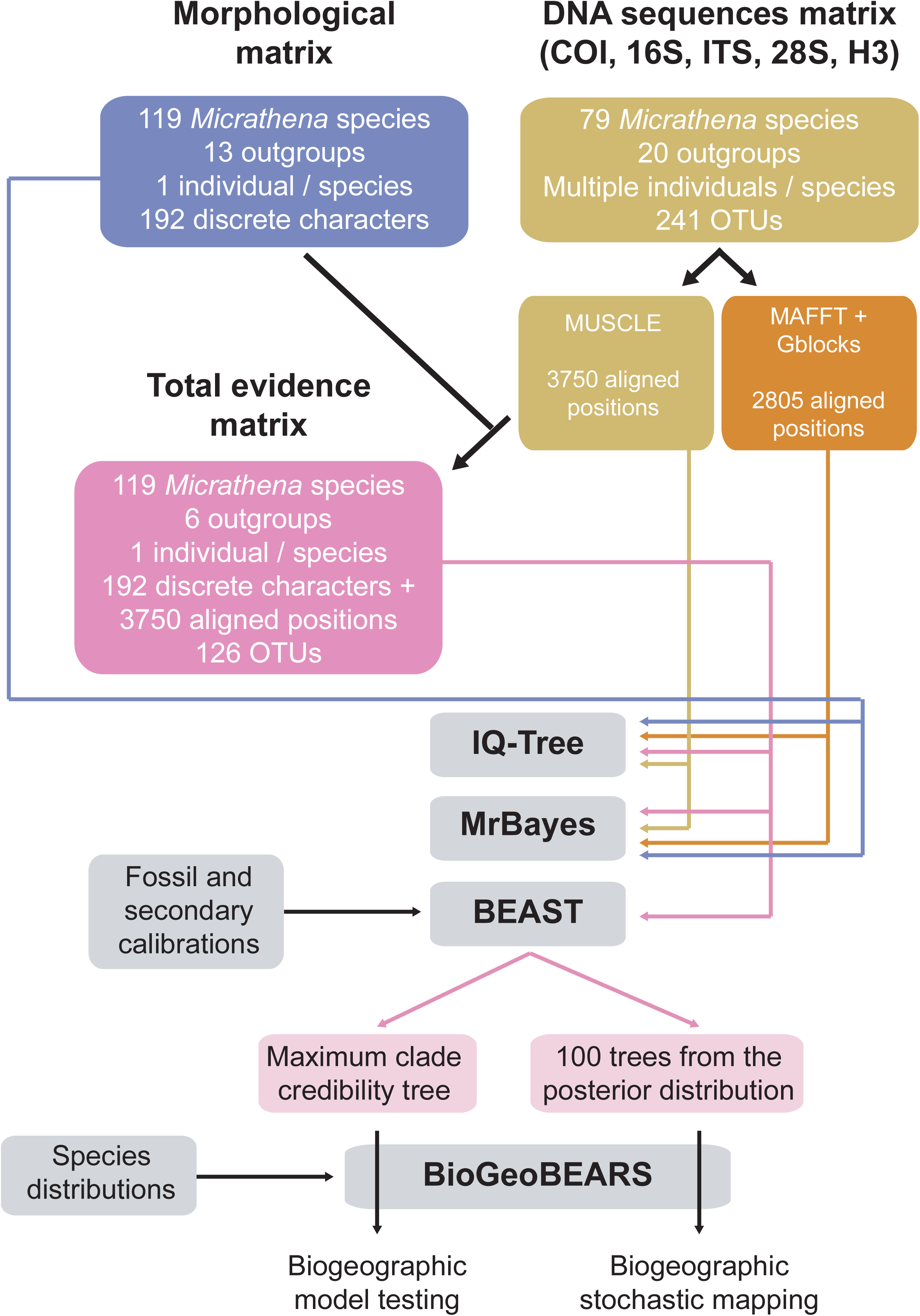
Summarized methodological pipeline used in this study.

**Figure S2.**
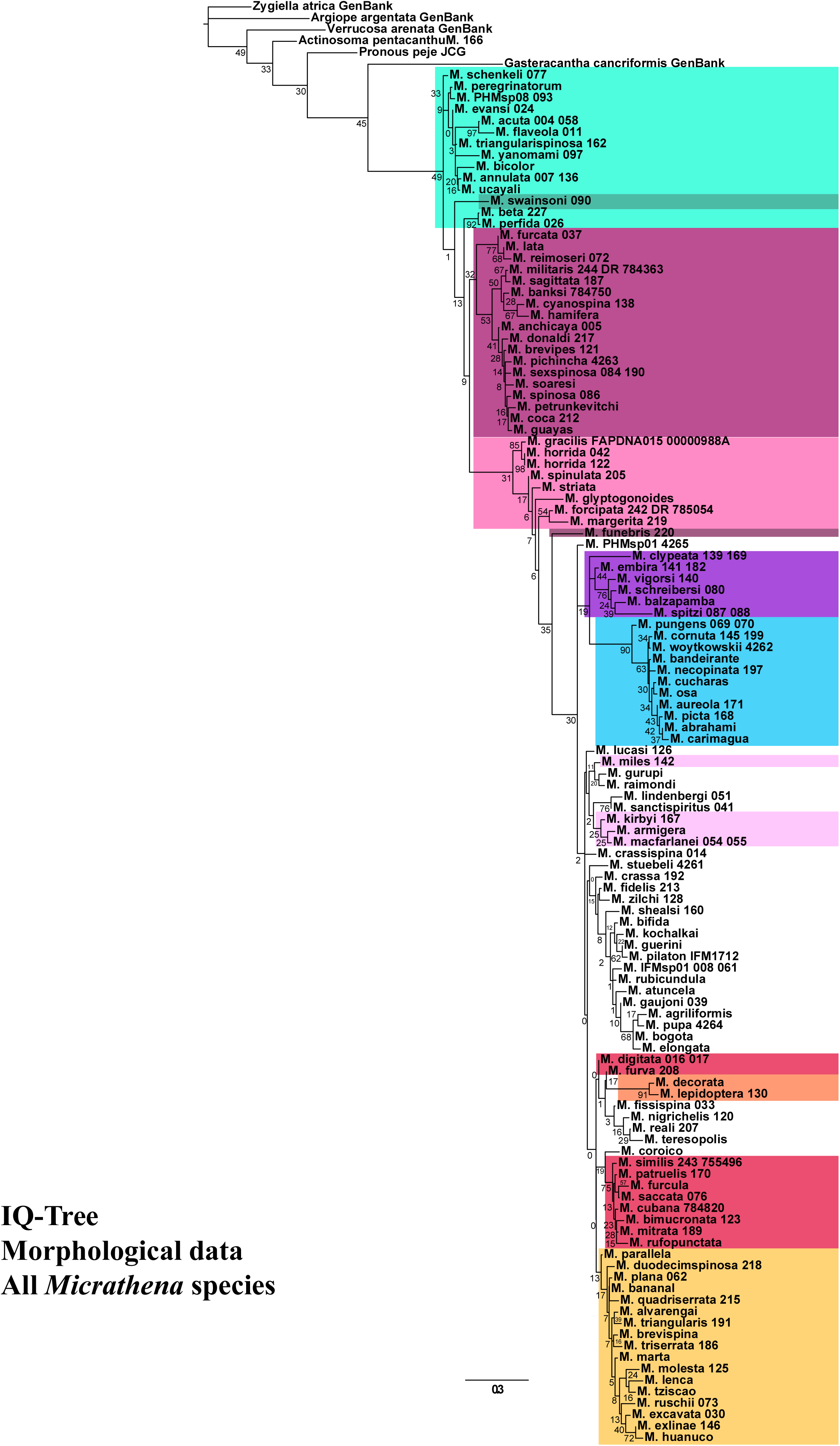
Maximum likelihood tree obtained by analyzing morphological data only under the MK+G4+F+ASC model in IQ-Tree. Scale bar represents the expected number of substitions per site. Numbers below nodes are bootstrap frequencies estimated from 100 pseudoreplicates. Colors correspond to those of Figure 2.

**Figure S3.**
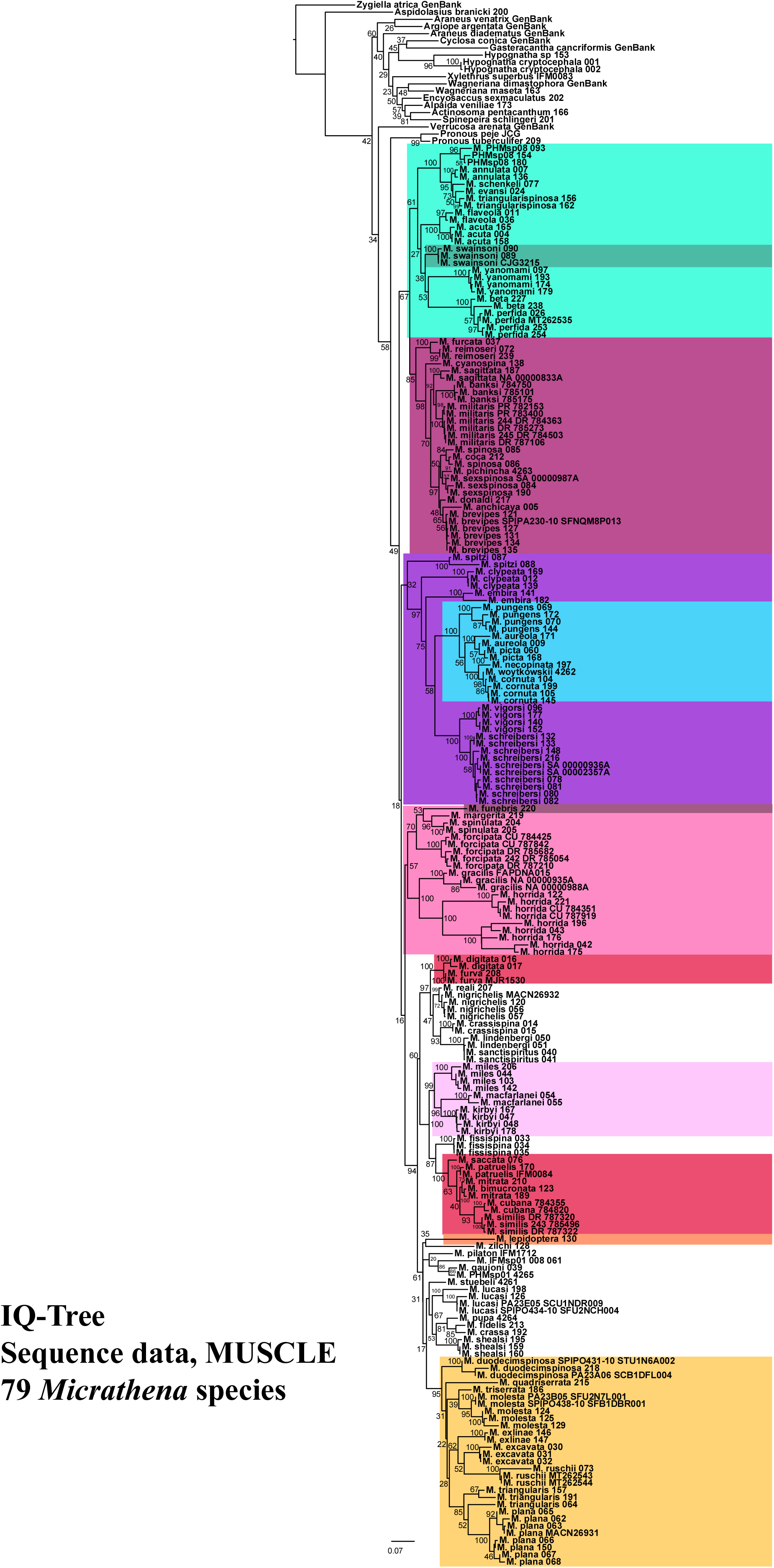
Maximum likelihood tree obtained by analyzing sequence data only (COI, 16S, ITS, 28S and H3, aligned with MUSCLE) in IQ-Tree. Scale bar represents the expected number of substitions per site. Numbers below nodes are bootstrap frequencies estimated from 100 pseudoreplicates. Colors correspond to those of Figure 2.

**Figure S4.**
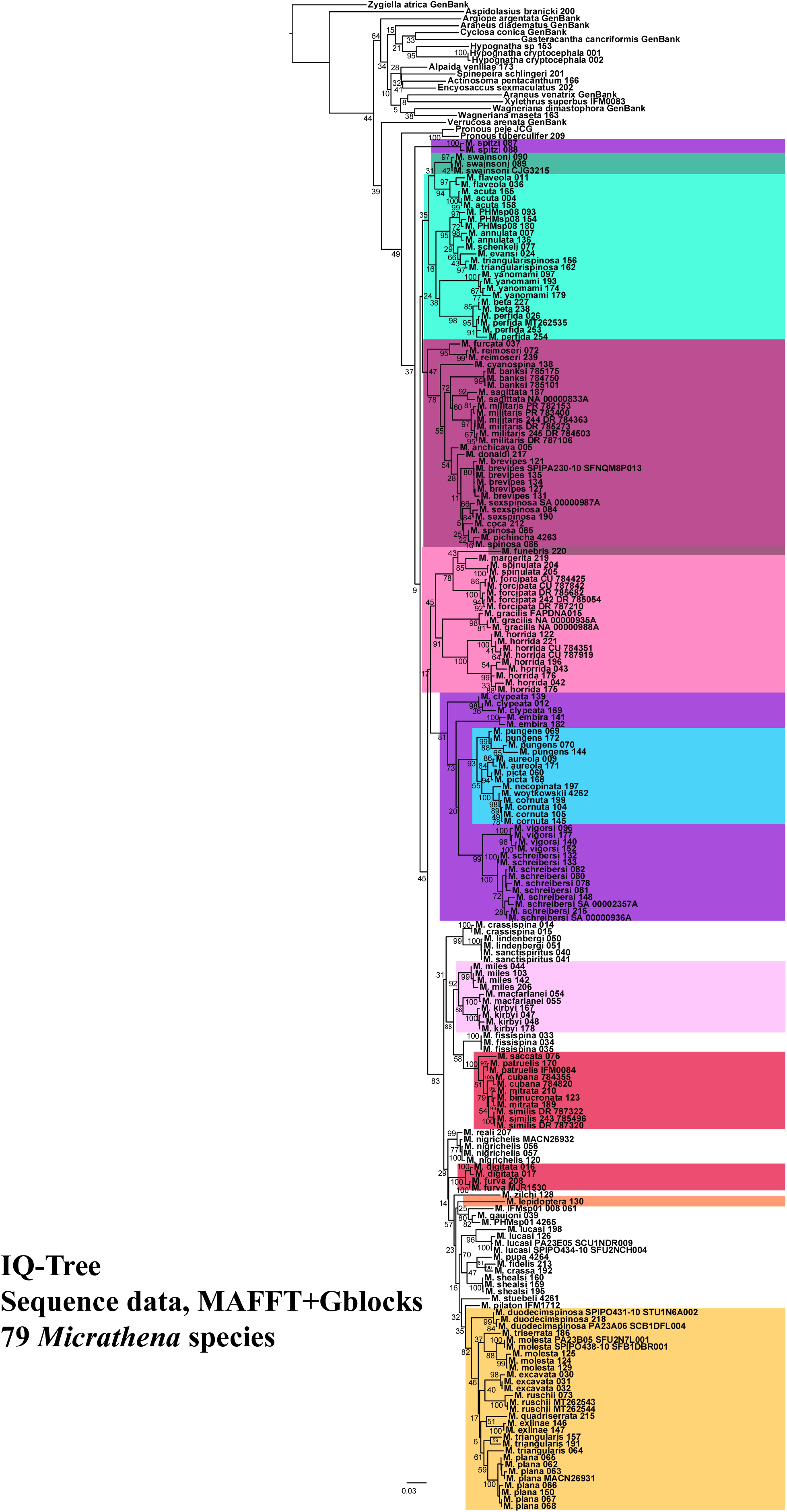
Maximum likelihood tree obtained by analyzing sequence data only (COI, 16S, ITS, 28S and H3, aligned with MAFFT and filtered with Gblocks) in IQ-Tree. Scale bar represents the expected number of substitions per site. Numbers below nodes are bootstrap frequencies estimated from 100 pseudoreplicates. Colors correspond to those of Figure 2.

**Figure S5.**
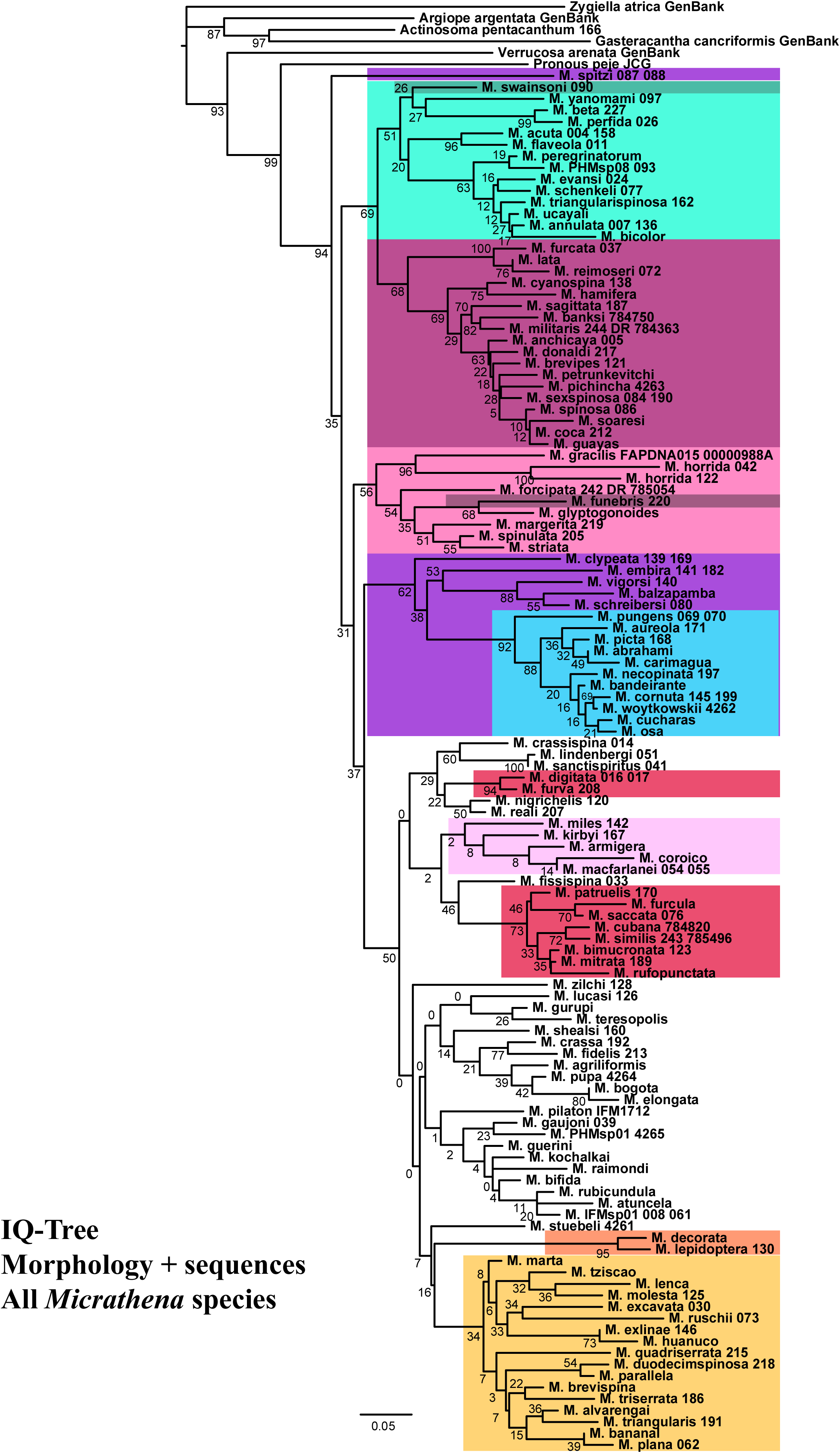
Maximum likelihood tree obtained by analyzing morphology and sequence data (COI, 16S, ITS, 28S and H3, aligned with MUSCLE) in IQ-Tree. Scale bar represents the expected number of substitions per site. Numbers below nodes are bootstrap frequencies estimated from 100 pseudoreplicates. Colors correspond to those of Figure 2.

**Figure S6.**
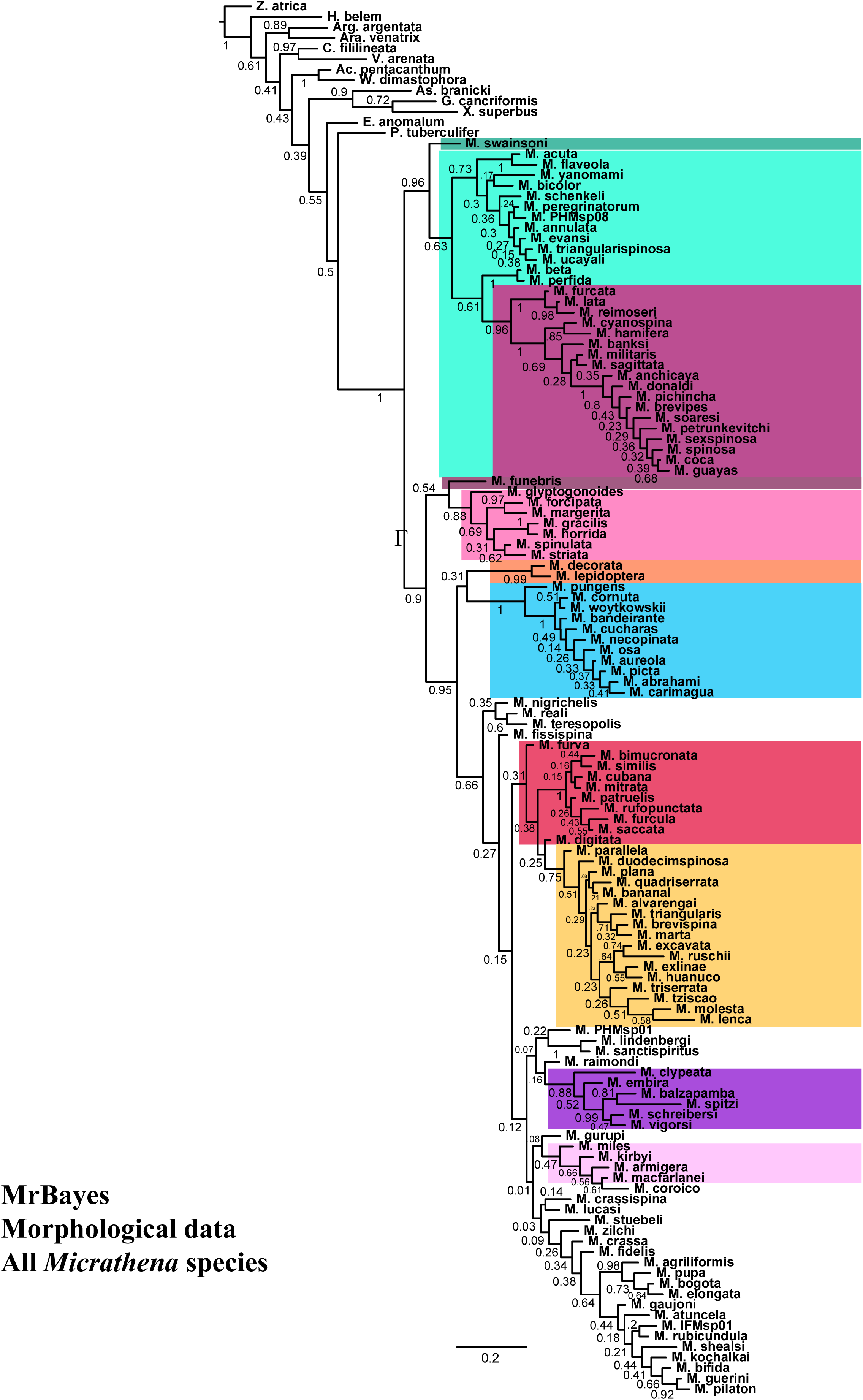
Majority-rule consensus of trees obtained by analyzing morphological data only under the Mkv + Γ model in MrBayes. Scale bar represents the expected number of substitions per site. Numbers below nodes are posterior probabilities. Colors correspond to those of Figure 2.

**Figure S7.**
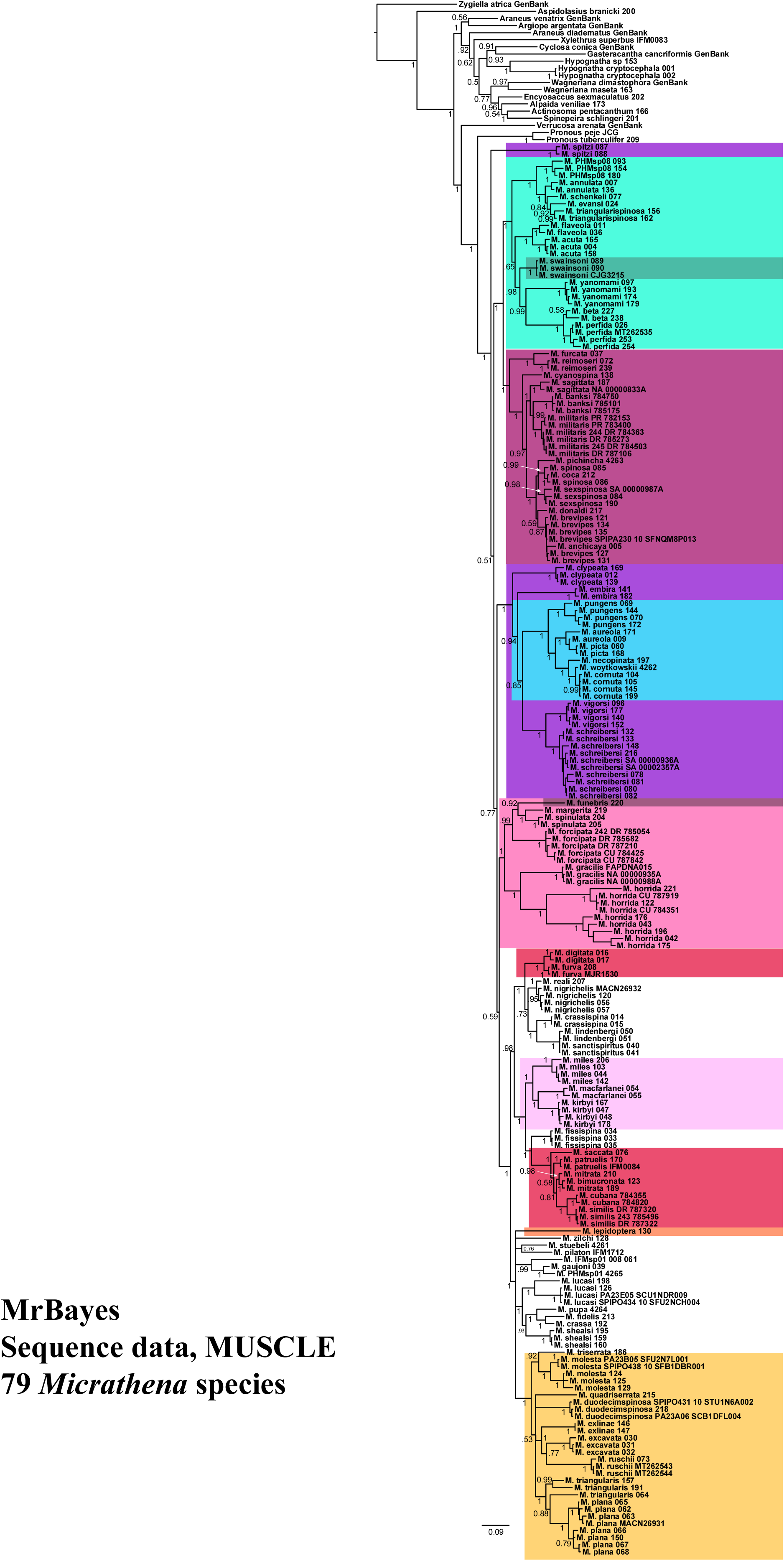
Majority-rule consensus of trees obtained by analyzing analyzing sequence data only (COI, 16S, ITS, 28S and H3, aligned with MUSCLE) in MrBayes. Scale bar represents the expected number of substitions per site. Numbers below nodes are posterior probabilities. Colors correspond to those of Figure 2.

**Figure S8.**
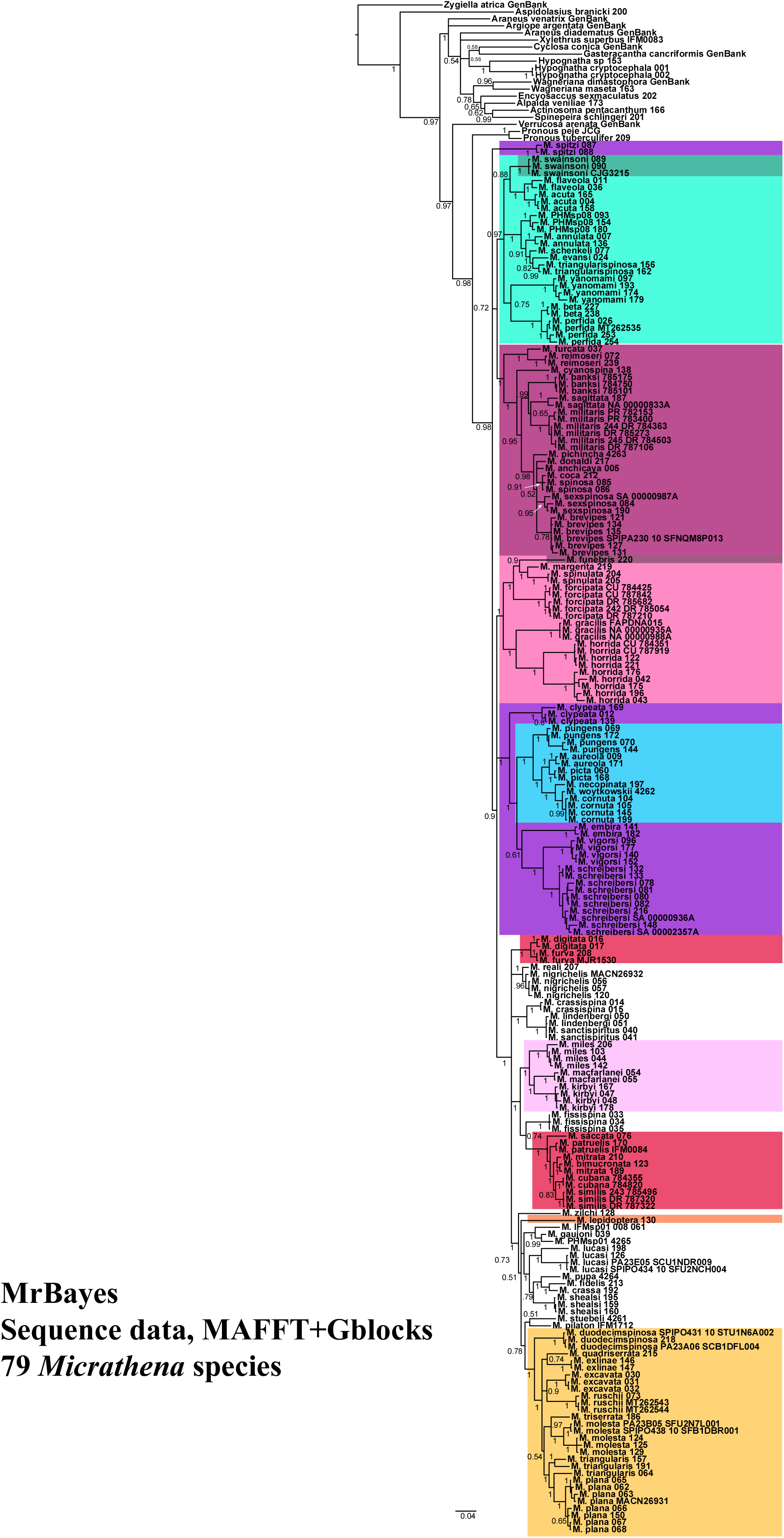
Majority-rule consensus of trees obtained by analyzing sequence data only (COI, 16S, ITS, 28S and H3, aligned with MAFFT and filtered with Gblocks) in MrBayes. Scale bar represents the expected number of substitions per site. Numbers below nodes are posterior probabilities. Colors correspond to those of Figure 2.

**Figure S9.**
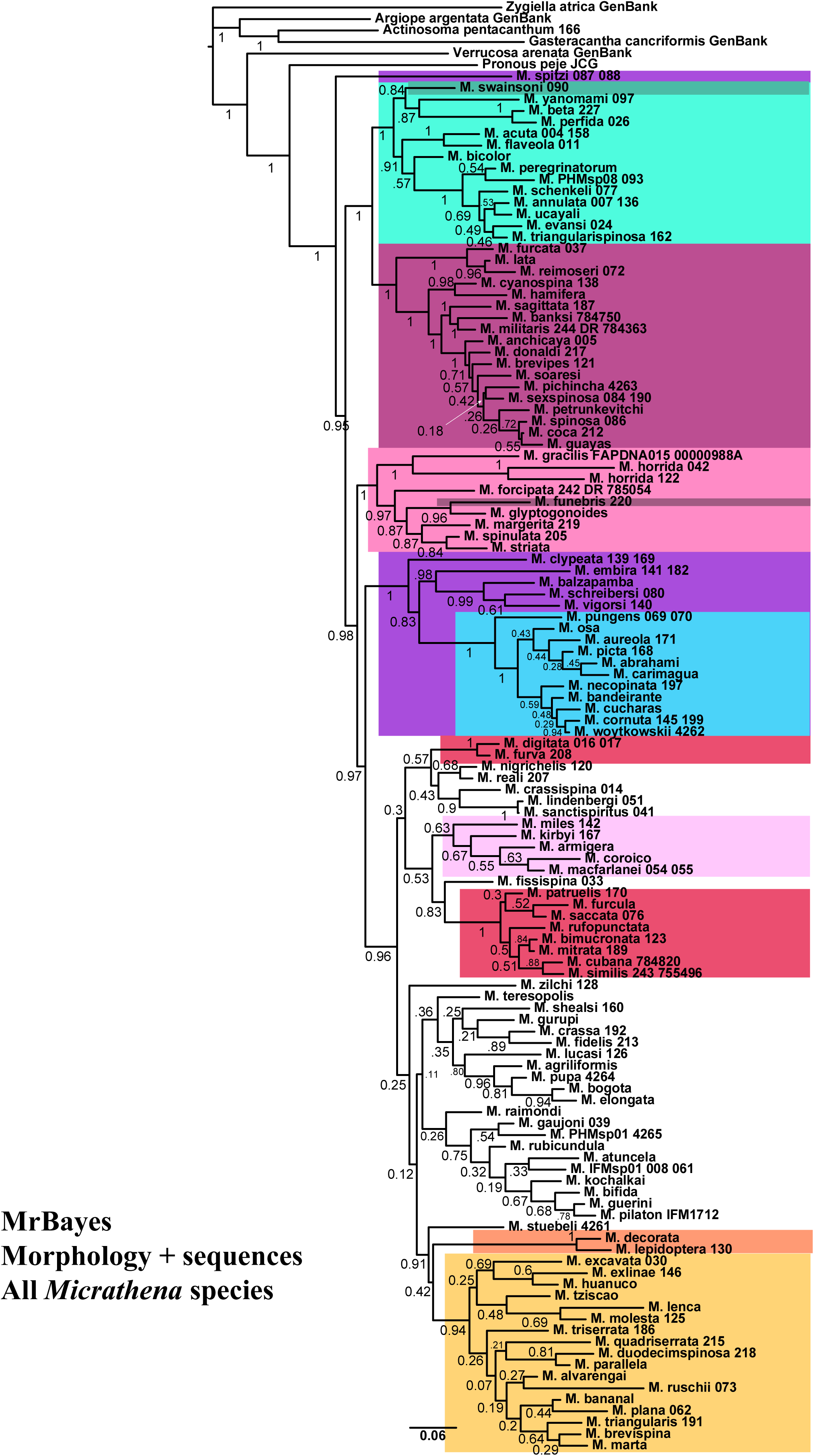
Majority-rule consensus of trees obtained by analyzing morphology and sequence data (COI, 16S, ITS, 28S and H3, aligned with MUSCLE) in MrBayes. Scale bar represents the expected number of substitions per site. Numbers below nodes are posterior probabilities. Colors correspond to those of Figure 2.

**Figure S10.**
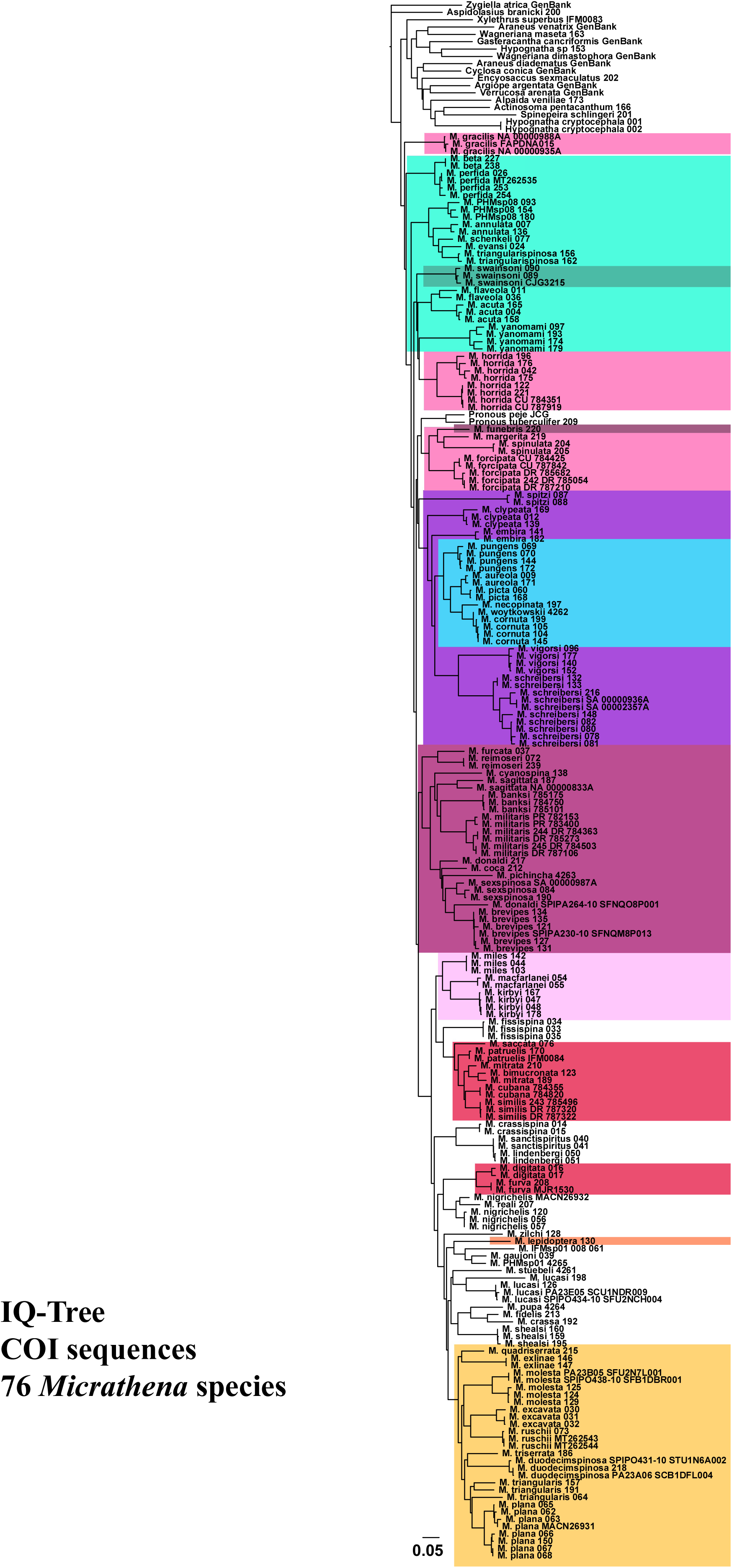
Maximum likelihood tree obtained by analyzing COI sequence data in IQ-Tree. Scale bar represents the expected number of substitions per site. Colors correspond to those of Figure 2.

**Figure S11.**
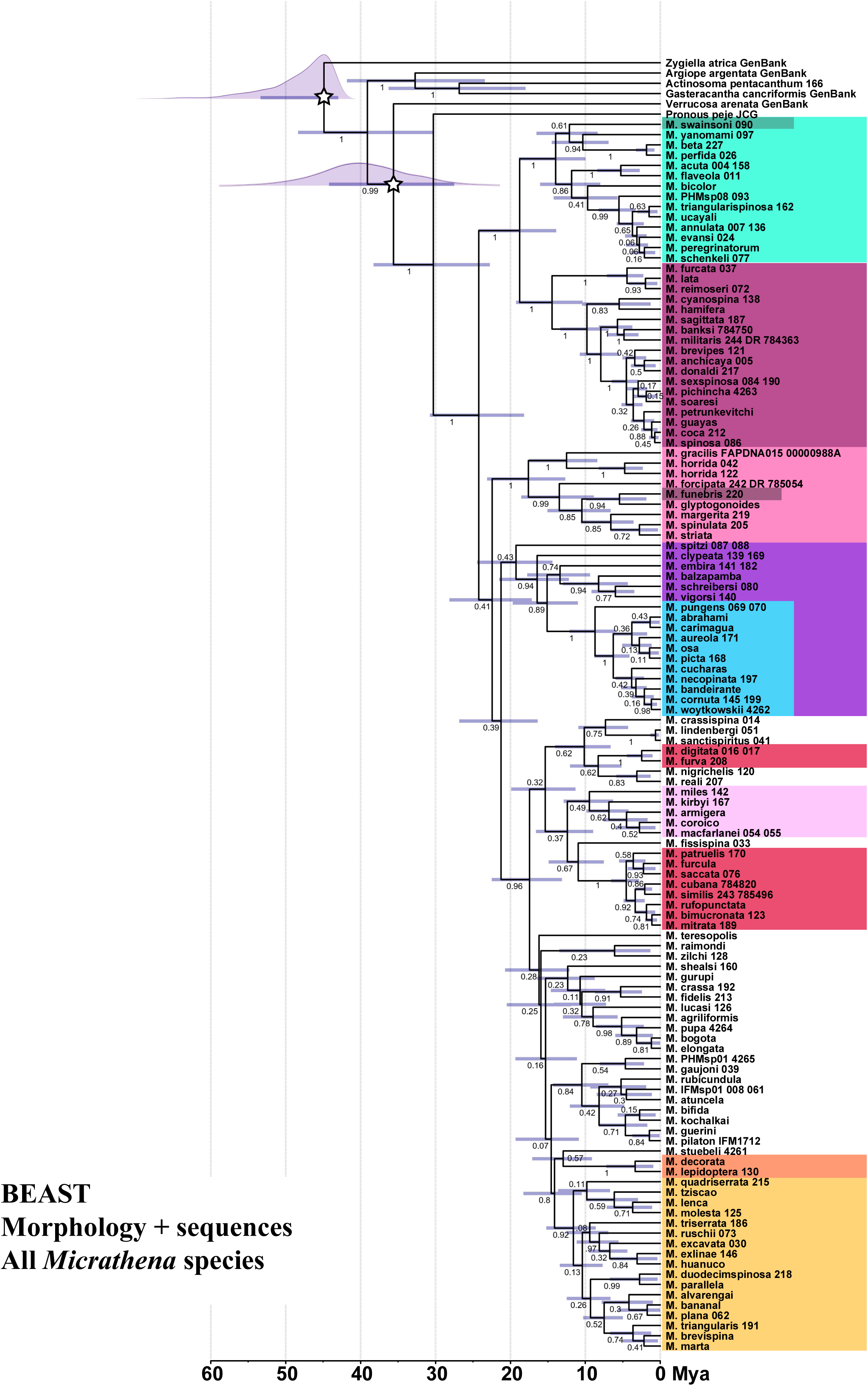
Maximum clade credibility tree obtained by analyzing morphology and sequence data (COI, 16S, ITS, 28S and H3, aligned with MUSCLE) in BEAST. Divergence times were estimated under a lognormal relaxed clock model. Stars mark the two calibrated nodes (see text for details); curves are the density of the ages of these nodes while sampling from the prior distribution only. Node bars are age 95% highest posterior density intervals. Numbers below nodes are posterior probabilities. Colors correspond to those of Figure 2.

**Figure S12.**
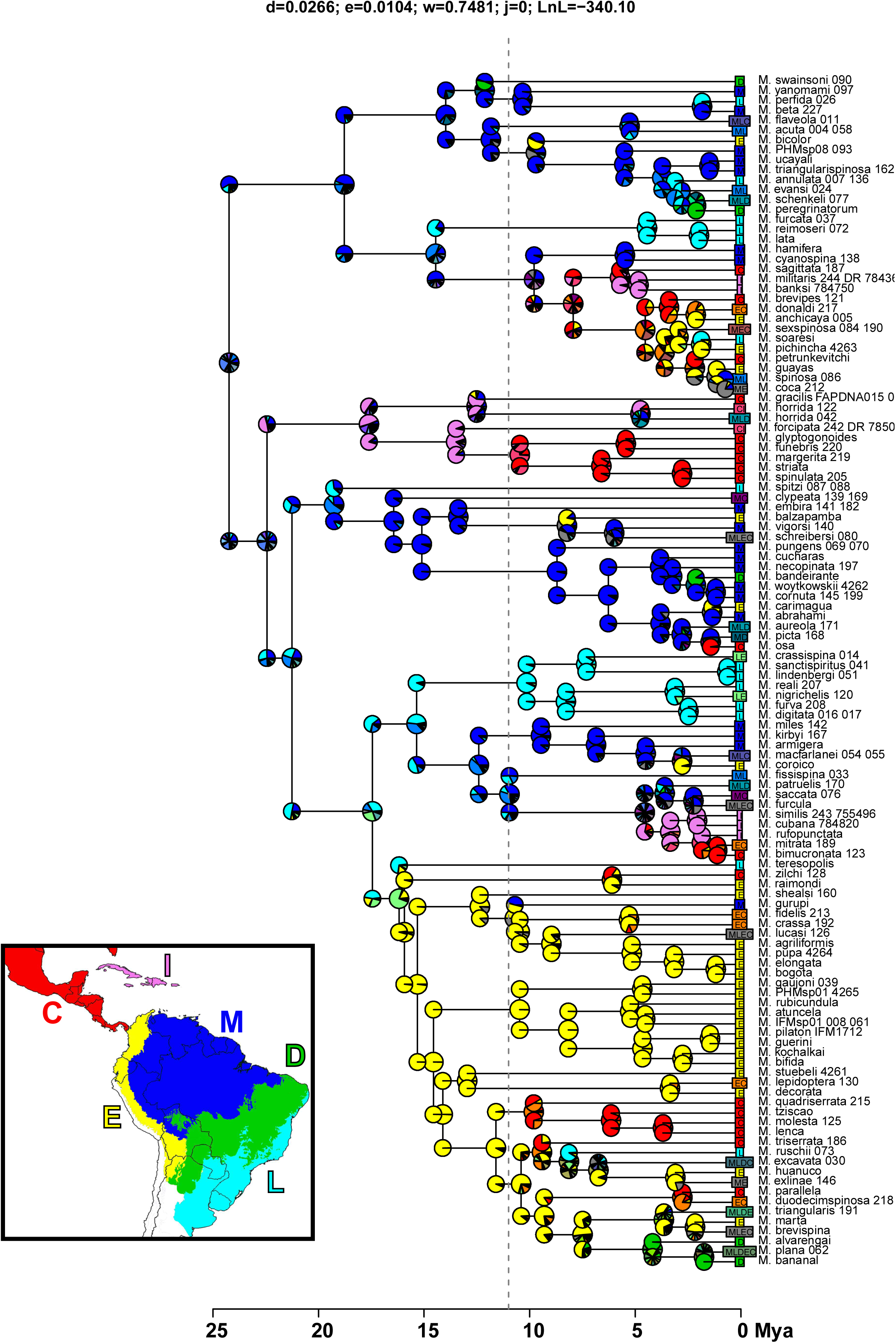
Ancestral ranges estimated for *Micrathena* under a time-stratified DEC model with Central America unavailable for occupation before 11 Mya.

**Figure S13.**
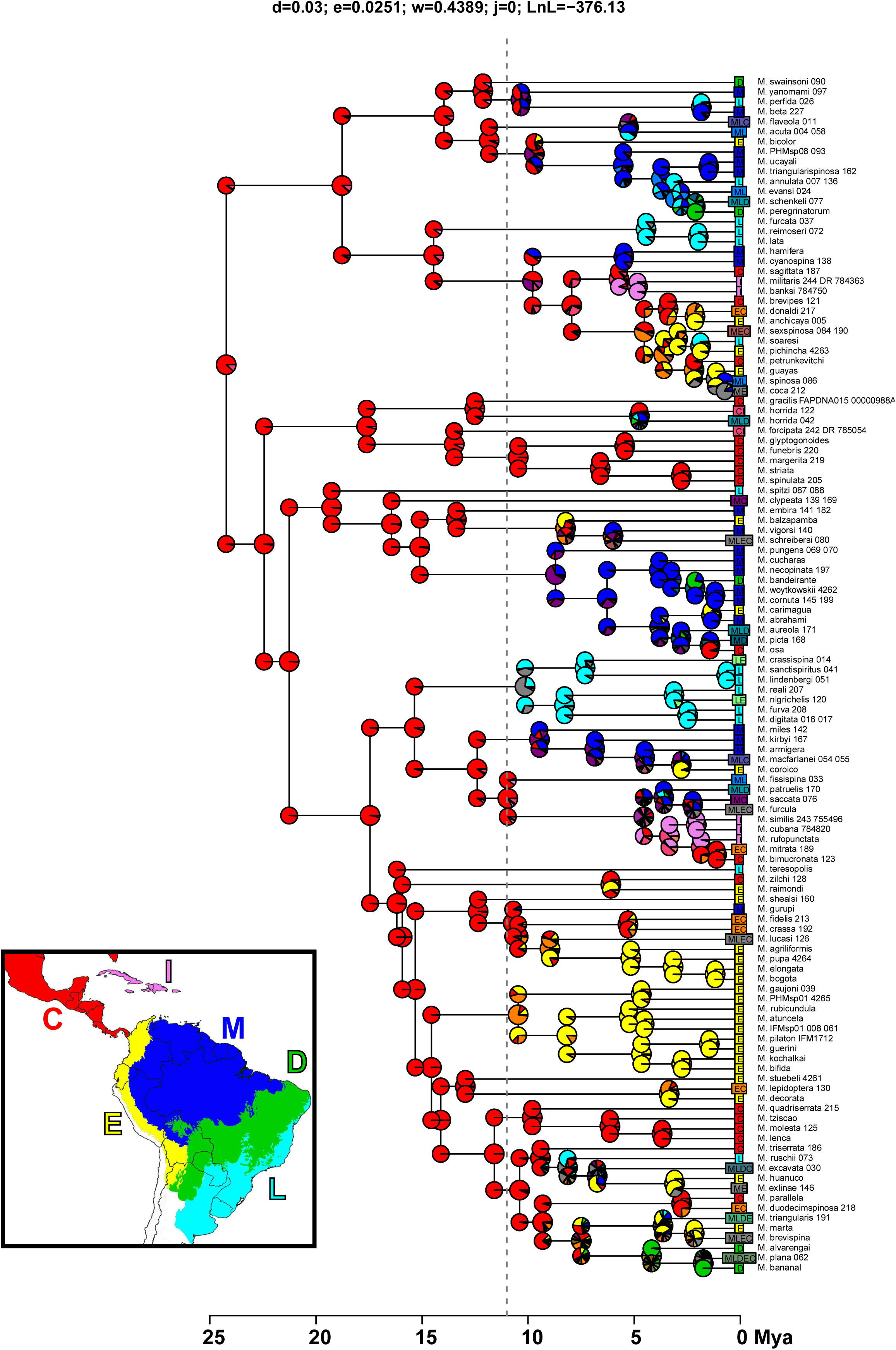
Ancestral ranges estimated for *Micrathena* under a time-stratified DEC model with South America (areas M, D, E and L) unavailable for occupation before 11 Mya.

**Figure S14.**
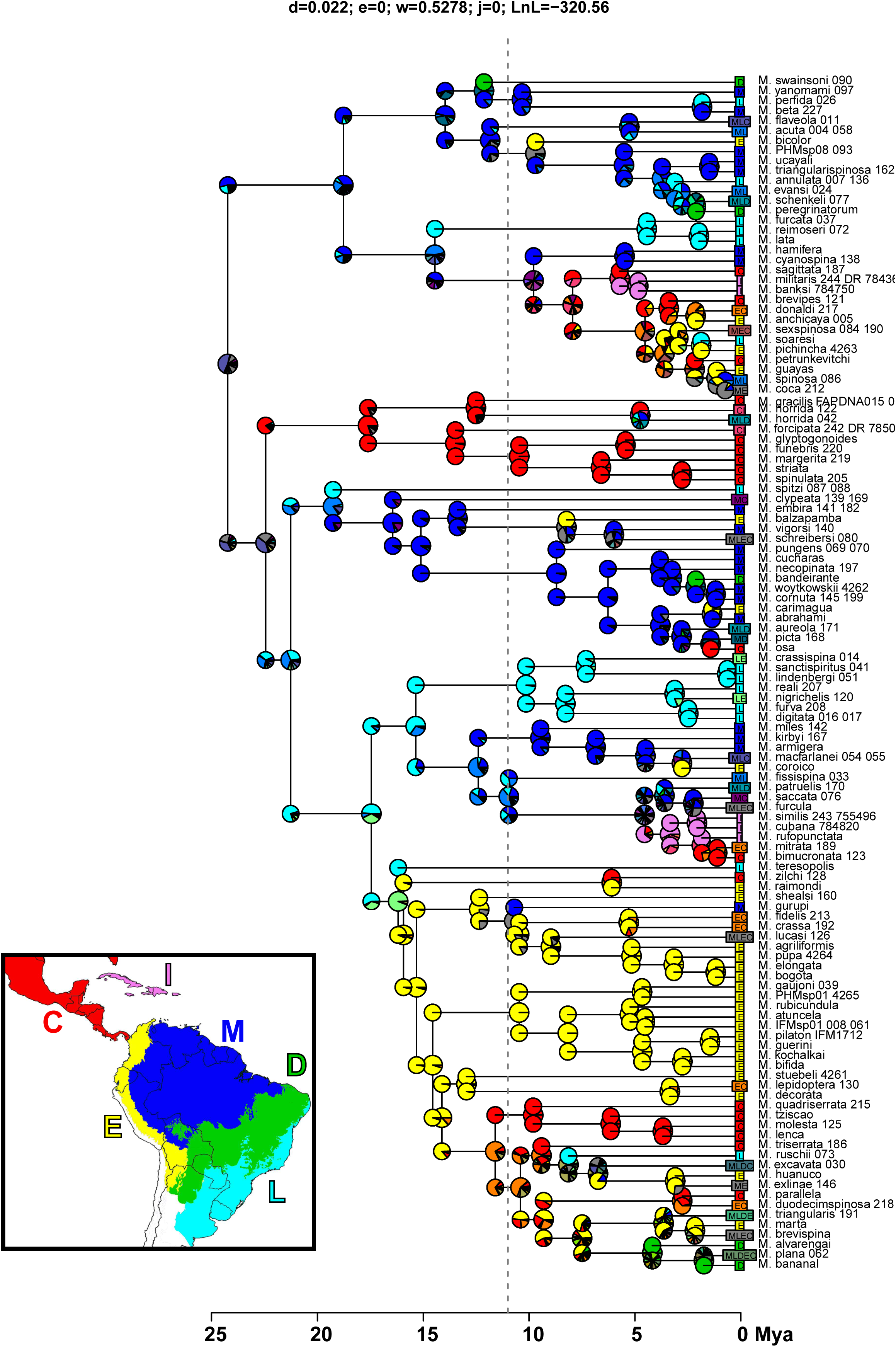
Ancestral ranges estimated for *Micrathena* under a time-stratified DEC model with dispersal probabilities between South and Central America reduced to 10% of the current values before 11 Mya.

**Figure S15.**
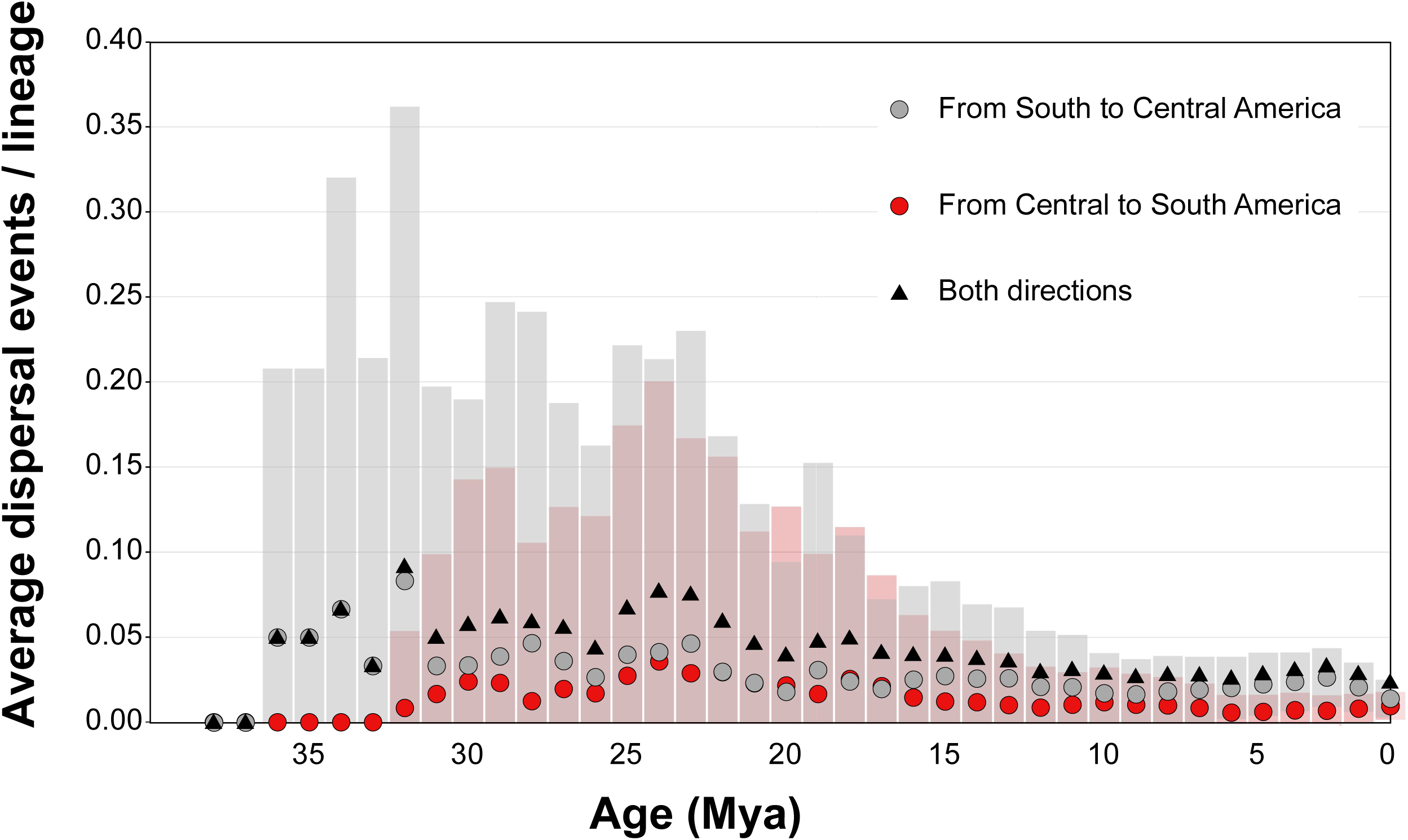
Rates of dispersal of *Micrathena* between South and Central America through time, corrected by the number of lineages in each time bin. The symbols are the dispersal events per lineage averaged over 1000 possible biogeographic histories (100 trees X 10 biogeographic stochastic maps each); bars are standard deviations. Rates of dispersal have not substantially increased after closure of the Isthmus of Panama.

## Notes

### Competing Interest Statement

The authors have declared no competing interest.

